# ARP-T1 is a ciliogenesis protein associated with a novel ciliopathy in inherited basal cell cancer, Bazex-Dupré-Christol Syndrome

**DOI:** 10.1101/2019.12.16.877498

**Authors:** Hyun-Sook Park, Eirini Papanastasi, Gabriela Blanchard, Elena Chiticariu, Daniel Bachmann, Markus Plomann, Fanny Morice-Picard, Pierre Vabres, Asma Smahi, Marcel Huber, Christine Pich, Daniel Hohl

## Abstract

Actin-Related Protein-Testis1 (ARP-T1)/ *ACTRT1* gene mutations cause the Bazex-Dupré-Christol Syndrome (BDCS) characterized by follicular atrophoderma, hypotrichosis and basal cell cancer. Here, we report an ARP-T1 interactome (PXD016557) involved in ciliogenesis, endosomal recycling and septin ring formation. Consequently, ARP-T1 localizes to the midbody during cytokinesis and the basal body of primary cilia in G_0_. Tissue samples from BDCS patients show reduced ciliary length with significant correlations of ARP-T1 expression levels, confirmed by *ACTRT1* knock down. We report that BDCS is a novel ciliopathy and the first case of a skin cancer ciliopathy, where ARP-T1 plays a critical role to prevent pathogenesis.

## INTRODUCTION

Basal cell cancer (BCC) of the skin is the most frequent human cancer. Bazex-Dupré-Christol Syndrome (BDCS) is an X-linked dominantly inherited syndrome form of BCC, without male-to-male transmission, affecting primarily the hair follicle (HF), resulting in the triad of hypotrichosis, follicular atrophoderma and BCC ^1–3^. Hypotrichosis and follicular atrophoderma develop around birth and BCC early in adulthood. BDCS less frequently presents with milia, hypohidrosis, facial pigmentation and trichoepithelioma, and thus combines developmental cutaneous anomalies with tumor predisposition, i.e. an ectodermal dysplasia with a hereditary tumor syndrome ^4,5^. Unlike most cases of BCC, <u>which are sporadic,</u> BDCS is an inherited syndromic form of BCC. The insertion mutation *ACTRT1* 547_548InsA, creating a shift of the reading frame, results in a truncated protein of 198 amino acids <u>leading to a loss of function of the protein. ARP-T1 was also found depleted in families with</u> germline mutations of non-coding sequences surrounding *ACTRT1* postulated to belong to enhancers transcribed as non-coding RNAs<u>. As these mutations segregate with the disease,</u> ARP-T1 <u>can be considered</u> as a tumor suppressor in BDCS ^6^. ARP-T1 was first isolated as a major acidic component of the cytoskeletal calyx from the head of bull spermatozoa. ARP-T1 is expressed specifically late in spermatid differentiation in testis, where it locates to the postacrosomal region and the centriole ^7^, establishing a link to the primary cilium (PC) of epidermal cells implicated in BCC development.

Cilia are microtubule-based organelles that are either motile as in sperm propel or nonmotile (primary) acting as sensory antenna, receiving signals from other cells nearby. The sensory capacity of PC is founded on the spatio-temporal localization of diverse receptor complexes such as PTCH, RTKs, TGFβR, NOTCH receptors, GPCRs including SMO, ion channels and extracellular matrix receptors along the cilium ^8^. Ciliary communication is often compromised in cancer ^9^, and faulty Hedgehog (HH) signaling implicated prototypically in medulloblastoma and BCC. Indeed, BCC cells are frequently ciliated, and activated HH signaling within PC is a key driver in BCC pathogenesis ^10^. BCCs generally show abnormal activation of the HH signaling pathway, ascribing its constitutive activation as a prerequisite for the tumor development ^11^. HH signaling in its “off” state is characterized by PTCH1 repressing SMO activity. PC have a dual role and are required for or may repress tumor formation ^12,13^, depending on the level of the molecular interaction ^9^.

PC assembly is a complex process emanating from the mother centriole with its appendages and differentiating upon cell cycle exit: the basal body, which nucleates the PC with apical CEP164 ^14^, a component of distal appendages, EHD1/EHD3 and SNAP29 ^15^ to initiate primary vesicle formation and ciliogenesis. Assembly of the PC includes recruitment of periciliary membranes, transition zone components and a machinery of intraflagellar transport proteins to form the ciliary vesicle and to set the base for formation of the microtubular ciliary axoneme ^15,16^. The PC is separated from the cytosol by the basal body and anchored by transition fibers/distal appendages in the ciliary pocket or periciliary plasma membrane (PM) ^17^. Transition fibers and transition zone with its Y-shaped linkers serve as a ciliary gate for the cilio-cytoplasmic transport sharing some functional similarities with the nuclear pore complex ^17,18^. Polarized membrane trafficking to the cilium and the vesicular transport machinery, which targets Golgi derived vesicles and apical recycling endosomes containing essential cargo, such as GPCRs, to the periciliary PM or the ciliary pocket depends on small GTPases, notably RAB8 and RAB11, Rabin 8 and the BBSome, a stable complex of seven proteins implicated in the ciliopathy of Bardet-Biedl Syndrome (BBS) ^8^.

Here, we demonstrate that ARP-T1 is a <u>basal body</u> protein and is involved in ciliogenesis by interacting with the ciliary machinery. <u>Mutations</u> in <u>*ACTRT1* or its enhancer elements,</u> as found in the tumor samples of BDCS patients<u>, as well as *ACTRT1* knockdown give</u> rise to the abnormally shortened cilia<u>, and this may be caused</u> by a disordered <u>diffusion barrier</u>. Ciliopathies encompass most human organ systems in eyes, nose, ears, organ placement, energy homeostasis, infertility, skeleton, reproductive system, brain, hydrocephalus, heart, chronic respiratory problems, kidney and liver ^19^. We report that BDCS is the first ciliopathy of epidermal development and cancer.

## RESULTS

### ARP-T1 is expressed in differentiated human keratinocytes and human epithelial cells

We examined the expression of ARP-T1 in NHEK (Fig. 1a,b) and HaCaT (Fig. 1c,d) cultured under differentiating conditions with high calcium up to seven days. The expression of ARP-T1 increased in a terminal differentiation dependent manner as indicated by the expression of Keratin 10 (K10) in differentiating keratinocytes at both mRNA (Fig. 1a,c) and protein levels (Fig. 1b,d). To investigate whether ARP-T1 expression was only induced in keratinocytes or whether it was related to differentiation, we also analyzed its expression in two differentiating retinal-pigmented epithelial (RPE) cells, ARPE-19 (Fig. 1e,f) and hTERT-RPE1 (Fig. 1g,h). These cells were differentiated under serum starvation for 35 days or 48 h respectively. <u>During RPE differentiation, ciliogenesis is known to increase ^14,20,21^ and IFT88 is a well-characterized protein involved in RPE ciliogenesis ^22,23^. We</u> assessed <u>RPE differentiation monitoring</u> with the expression of the ciliary protein IFT88 (Fig. 1f,h<u>), and with the increased number of ciliated cells (Extended data Fig. 1a</u>). Similar to keratinocytes, differentiated RPE cells exhibited an increased ARP-T1 expression after differentiation, at both mRNA (Fig. 1e,g) and protein levels (Fig. 1f,h, <u>quantified in Extended data Fig. 1b,c</u>).

**Figure 1.**
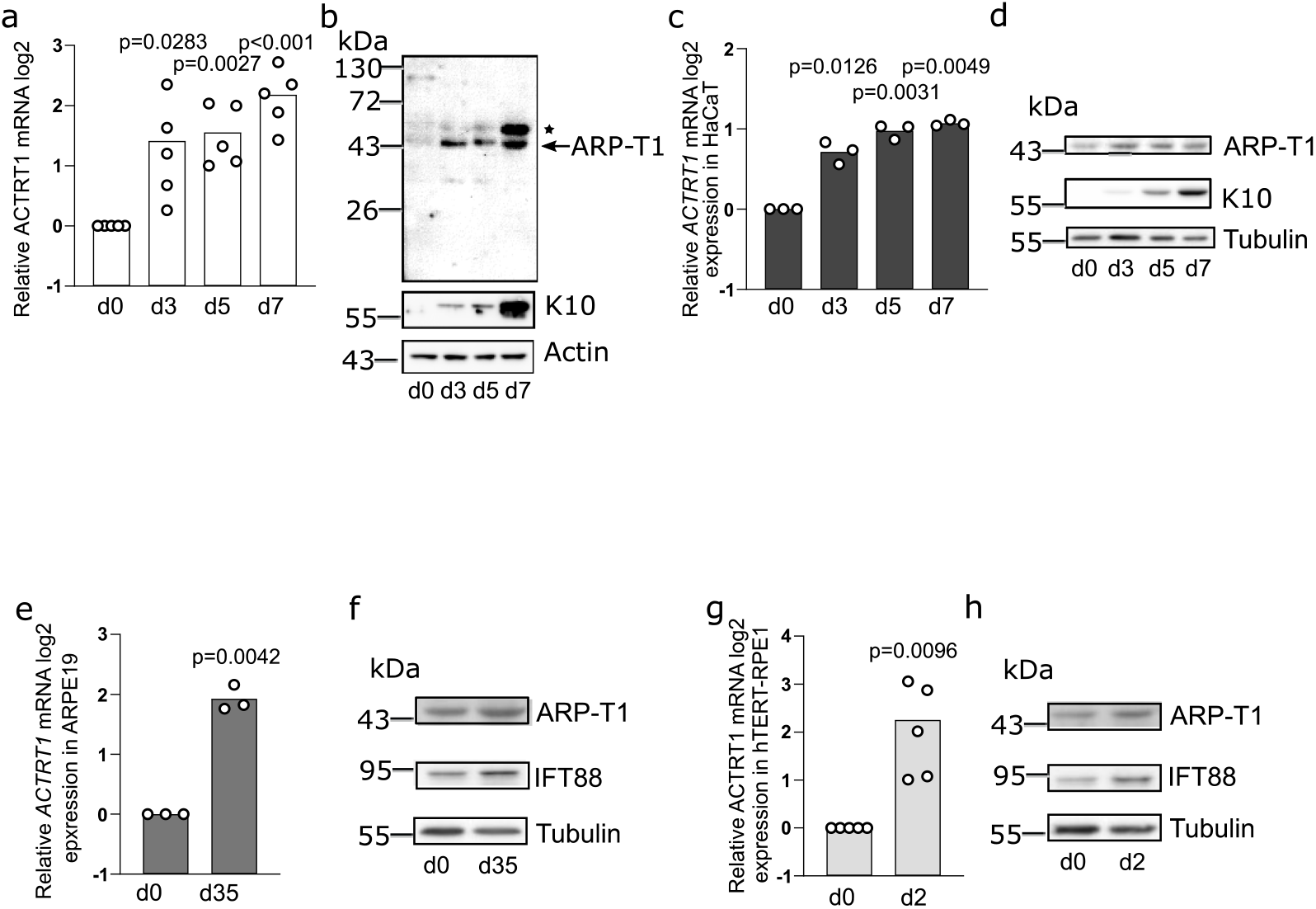
ARP-T1 is expressed during epidermal and epithelial differentiation. **a**,**c**,**e**,**g**, mRNA expression of *ACTRT1* during differentiation of keratinocytes, NHEK (**a**, N=5) and HaCaT (**c**, N=3), and epithelial cells, ARPE19 (**e**, N=3) and hTERT-RPE1 (**g**, N=5). Data are presented as means of the fold change compared to the value of undifferentiated samples. <u>Each open circle represents one independent experiment.</u> **b**,**d**,**f**,**h**, <u>Representative images of</u> ARP-T1 expression during differentiation of keratinocytes, NHEK (**b**) and HaCaT (**d**), and epithelial cells, ARPE19 (**f**) and hTERT-RPE1 (**h**). ARP-T1 was detected using guinea pig anti-ARP-T1 antisera (**b**) or mouse antibody (**d**,**f**,**h**). * indicates polymers of ARP-T1 confirmed by mass spectrometry analysis. Keratin 10 and IFT88 were used as markers of cell differentiation in keratinocytes and epithelial cells respectively, actin and tubulin as loading controls.

### *ACTRT1* <u>mRNA</u> is regulated by non-canonical hedgehog pathway and protein kinase C

ARP-T1 was previously reported to inhibit *GLI1* expression and be involved in HH signaling ^6^. We treated HH responsive hTERT-RPE1 cells with Smoothened (SMO) agonist (SAG), which binds to SMO and activates the HH pathway, and measured the expression of *ACTRT1*. *GLI1* and *PTCH1* mRNA expression was used as controls of HH pathway activation. *GLI1* and *PTCH1* mRNA responded well upon SAG treatment in proliferating and serum-starved cells (Fig. 2a,b respectively).The expression of *ACTRT1* mRNA <u>and ARP-T1 protein</u> was increased by SAG treatment in proliferating (Fig. 2a) and differentiating (Fig. 2b) conditions. We used additional SMO activator, purmorphamine, to confirm the results with SAG treatment, under differentiating condition. In this case, we also used vismodegib (SMO inhibitor used for the treatment of BCC) and both purmorphamine and vismodegib, to understand better the regulation of ARP-T1 by the HH pathway<u>. *ACTRT1*,</u> *GLI1* and *PTCH1* mRNA increased about 2 fold upon purmorphamine treatment similar as the treatment with SAG (Fig. 2c-e). *GLI1* and *PTCH1* expression decreased upon both purmorphamine and vismodegib treatment as expected (Fig. 2d,e), but not the expression of *ACTRT1* indicating that *ACTRT1* mRNA expression is regulated by a non-canonical HH pathway<u>, although ARP-T1 protein expression was similar to the basal level in this condition</u> (Fig. 2c).

**Figure 2.**
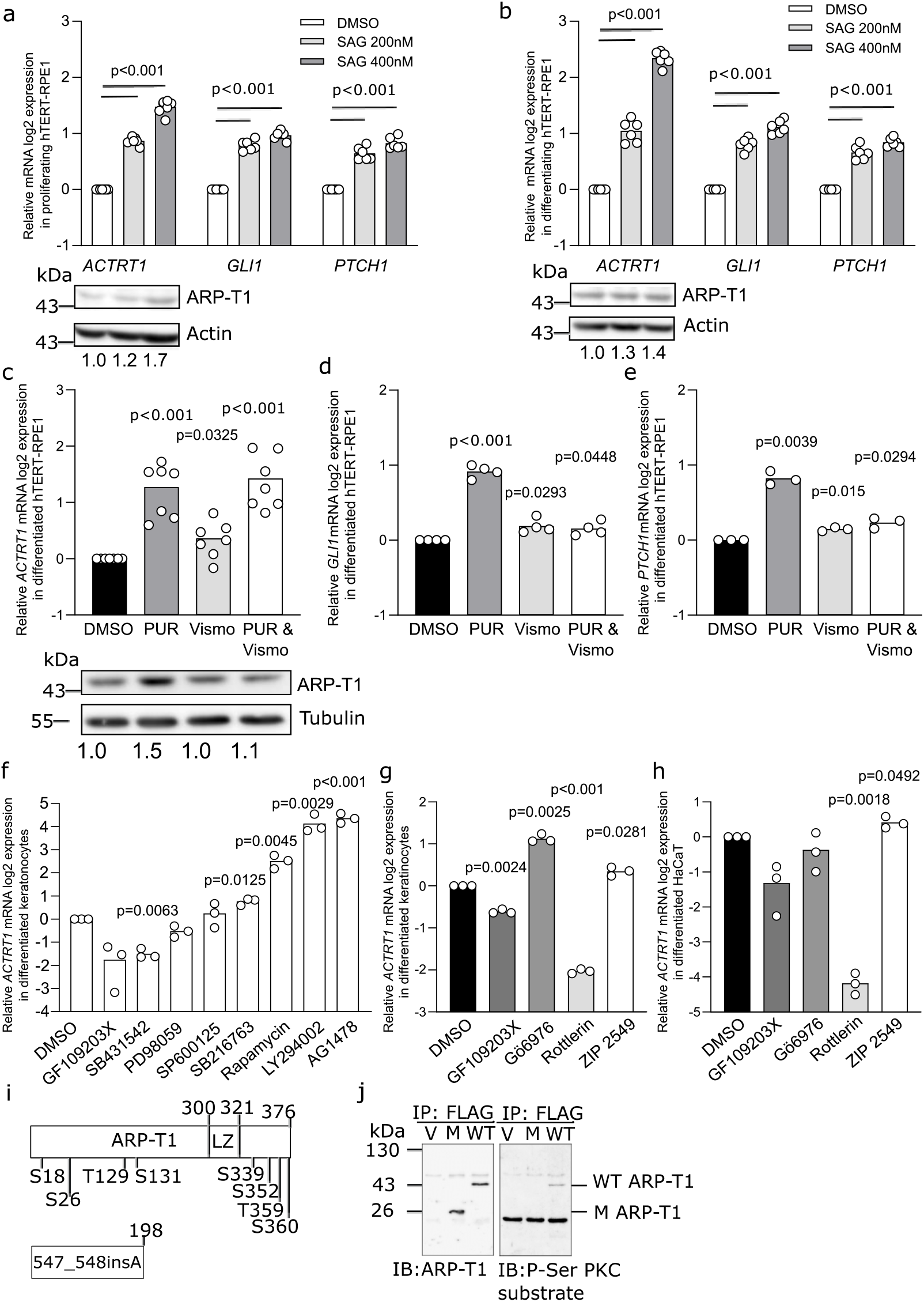
*ACTRT1* is regulated by non-canonical Hedgehog signaling pathway and by Protein Kinase C delta. **a**,**b**, Relative *ACTRT1*, *GLI1* and *PTCH1* mRNA expression <u>(**a,b**, N=6) and ARP-T1 expression (**a**, N=4; **b**, N=3)</u> upon treatment with SAG in hTERT-RPE1 cells under proliferative (**a**) and differentiating (**b**) conditions. <u>Actin is used as loading control for ARP-T1 expression. Numbers under the blots present the fold change expression of ARP-T1 compared to the vehicle (DMSO) treatment.</u> **c**-**e** Relative *ACTRT1* (**c**, N=7), *GLI1* (**d**, N=4), *PTCH1* (**e**, N=3) mRNA expression <u>and ARP-T1 expression (**c**, N=4)</u> upon treatment with purmorphamine <u>(PUR)</u> and/or vismodegib <u>(VISMO)</u> in differentiated hTERT-RPE1 cells. <u>Tubulin is used as loading control for ARP-T1 expression. Numbers under the blots present the fold change expression of ARP-T1 compared to the vehicle treatment.</u> **f**,**g**, Relative *ACTRT1* mRNA expression upon treatment with different protein kinase inhibitors (**f**) or with PKC inhibitors (**g**) in differentiated NHEK (N=3). **h**, Relative *ACTRT1* mRNA expression upon treatment with PKC inhibitors in differentiated HaCaT cells (N=3). <u>**a-h**, Data are presented as means of the fold change compared to the value of vehicle treated samples. Each open circle represents one independent experiment.</u> **i**, Schematic representation of ARP-T1 and ARP-T1 mutant (547_548insA) with predicted phosphorylation sites. **j**, ARP-T1 and Phospho-Serine (P-Ser) PKC expression in NHEK transduced with lentiviral vectors, empty vector (V), *ACTRT1* mutant (M) and *ACTRT1* WT (WT), after immunoprecipitated (IP) with anti-FLAG monoclonal antibody M2-conjugated agarose.

A cascade of phosphorylation is known to be involved in HH signaling pathways, e.g. PKC, GSK-3β, S6K, PKA, PI3K, aPKC-ι/λ ^8^. To investigate the HH pathway in which ARP-T1 plays a role in BCC onset, differentiated NHEK, in which ARP-T1 is highly expressed, were treated with different kinase inhibitors. The cells treated with GF109203X known as PKC and PKA inhibitor showed significantly decreased expression of *ACTRT1* mRNA, while cells treated with rapamycin (mTOR/S6K inhibitor), LY294002 (PI3K inhibitor) and AG1478 (epidermal growth factor tyrosine kinase inhibitor) showed increased expression of *ACTRT1* mRNA (Fig. 2f). We further narrowed down to treat NHEK and HaCaT cells with Gö6976 (PKC alpha/beta inhibitor), Rottlerin (PKC delta inhibitor) and ZIP 2549 (atypical PKC inhibitor). We found that Rottlerin inhibits the expression of *ACTRT1* mRNA in both NHEK and HaCaT cells, indicating that *ACTRT1* mRNA expression is regulated by PKC delta (Fig. 2g,h). Two different programs (PhosphositePlus and Swiss Institute of Bioinformatics) predicted several putative phosphorylation sites in ARP-T1 WT and the mutant protein (from *ACTRT1* 547_548insA mutation) (Fig. 2i). One of the predicted phosphorylation sites in C-terminus at S352 position showed similar consensus phosphorylation site specificity with PKC (www.kinexus.ca). We transduced *ACTRT1* (WT), *ACTRT1*547_548insA (M) and vector control (V) tagged with DDK (same sequence as FLAG) into differentiated NHEK and immunoprecipitated using anti-FLAG M2 affinity gel. We used the Phospho-(Ser) PKC substrate antiserum, which recognizes proteins only when phosphorylated at serine residues surrounded by arginine or lysine at the −2 and +2 positions and a hydrophobic residue at the +1 position (ARP-T1 S352 : MSS*FKQ). This antiserum recognized ARP-T1 WT but not ARP-T1 mutant suggesting that ARP-T1 is indeed phosphorylated by PKC and stable while ARP-T1 mutant, which lack the phosphorylation site, is not phosphorylated by PKC (Fig. 2j).

### ARP-T1 interacts with proteins <u>involved</u> in <u>ciliogenesis</u>

To understand the molecular pathways exploited by ARP-T1, we performed co-immunoprecipitation followed by mass spectrometry (MS) analysis to identify proteins interacting with ARP-T1. We transduced *ACTRT1* (WT), *ACTRT1* 547_548insA (M) and vector control (V) tagged with DDK into the cells and immunoprecipitated using anti-FLAG M2 beads. We found putative interacting proteins in HeLa cells (Table 1, Fig. 3a), differentiated hTERT-RPE1 cells (Table 1, Fig. 3b), and NHEK (Table 1). ARP-T1 interacts in all cells with proteins involved in PC structure. We confirmed these interactions by immunoblot analysis with acetylated-tubulin, TCP8 (one of 8 subunits of chaperonin-containing T-complex), HSC70, BAG2, gamma-tubulin, EHD4, septin 2 and septin 9 antisera (Fig. 3a,b). While chaperones are often identified to some extent in MS analysis of precipitates with overexpressed proteins, the high number of spectral hits (Table 1) and the location of septins and EHD4 at ciliary basal body led us to observe the localization of ARP-T1 in ciliated hTERT-RPE1 cells.

**Table 1.**
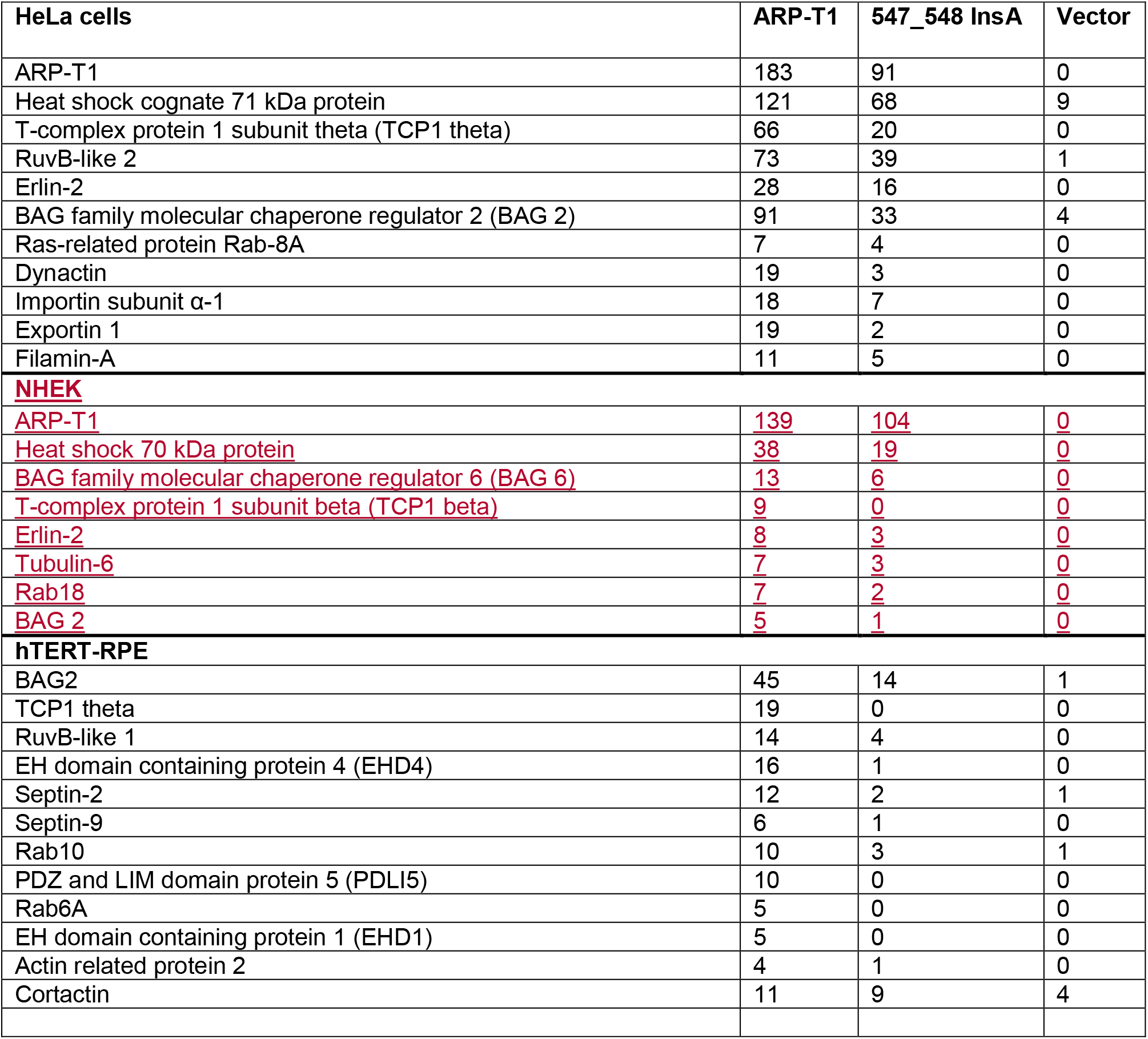

| HeLa cells | ARP-T1 | 547_548 InsA | Vector |
| --- | --- | --- | --- |
| ARP-T1 | 183 | 91 | 0 |
| Heat shock cognate 71 kDa protein | 121 | 68 | 9 |
| T-complex protein 1 subunit theta (TCP1 theta) | 66 | 20 | 0 |
| RuvB-like 2 | 73 | 39 | 1 |
| Erlin-2 | 28 | 16 | 0 |
| BAG family molecular chaperone regulator 2 (BAG 2) | 91 | 33 | 4 |
| Ras-related protein Rab-8A | 7 | 4 | 0 |
| Dynactin | 19 | 3 | 0 |
| Importin subunit α-1 | 18 | 7 | 0 |
| Exportin 1 | 19 | 2 | 0 |
| Filamin-A | 11 | 5 | 0 |
| **NHEK** |  |  |  |
| <u>ARP-T1</u> | <u>139</u> | <u>104</u> | <u>0</u> |
| <u>Heat shock 70 kDa protein</u> | <u>38</u> | <u>19</u> | <u>0</u> |
| <u>BAG family molecular chaperone regulator 6 (BAG 6)</u> | <u>13</u> | <u>6</u> | <u>0</u> |
| <u>T-complex protein 1 subunit beta (TCP1 beta)</u> | <u>9</u> | <u>0</u> | <u>0</u> |
| <u>Erlin-2</u> | <u>8</u> | <u>3</u> | <u>0</u> |
| <u>Tubulin-6</u> | <u>7</u> | <u>3</u> | <u>0</u> |
| <u>Rab18</u> | <u>7</u> | <u>2</u> | <u>0</u> |
| <u>BAG 2</u> | <u>5</u> | <u>1</u> | <u>0</u> |
| **hTERT-RPE** |  |  |  |
| BAG2 | 45 | 14 | 1 |
| TCP1 theta | 19 | 0 | 0 |
| RuvB-like 1 | 14 | 4 | 0 |
| EH domain containing protein 4 (EHD4) | 16 | 1 | 0 |
| Septin-2 | 12 | 2 | 1 |
| Septin-9 | 6 | 1 | 0 |
| Rab10 | 10 | 3 | 1 |
| PDZ and LIM domain protein 5 (PDLI5) | 10 | 0 | 0 |
| Rab6A | 5 | 0 | 0 |
| EH domain containing protein 1 (EHD1) | 5 | 0 | 0 |
| Actin related protein 2 | 4 | 1 | 0 |
| Cortactin | 11 | 9 | 4 |

**Figure 3.**
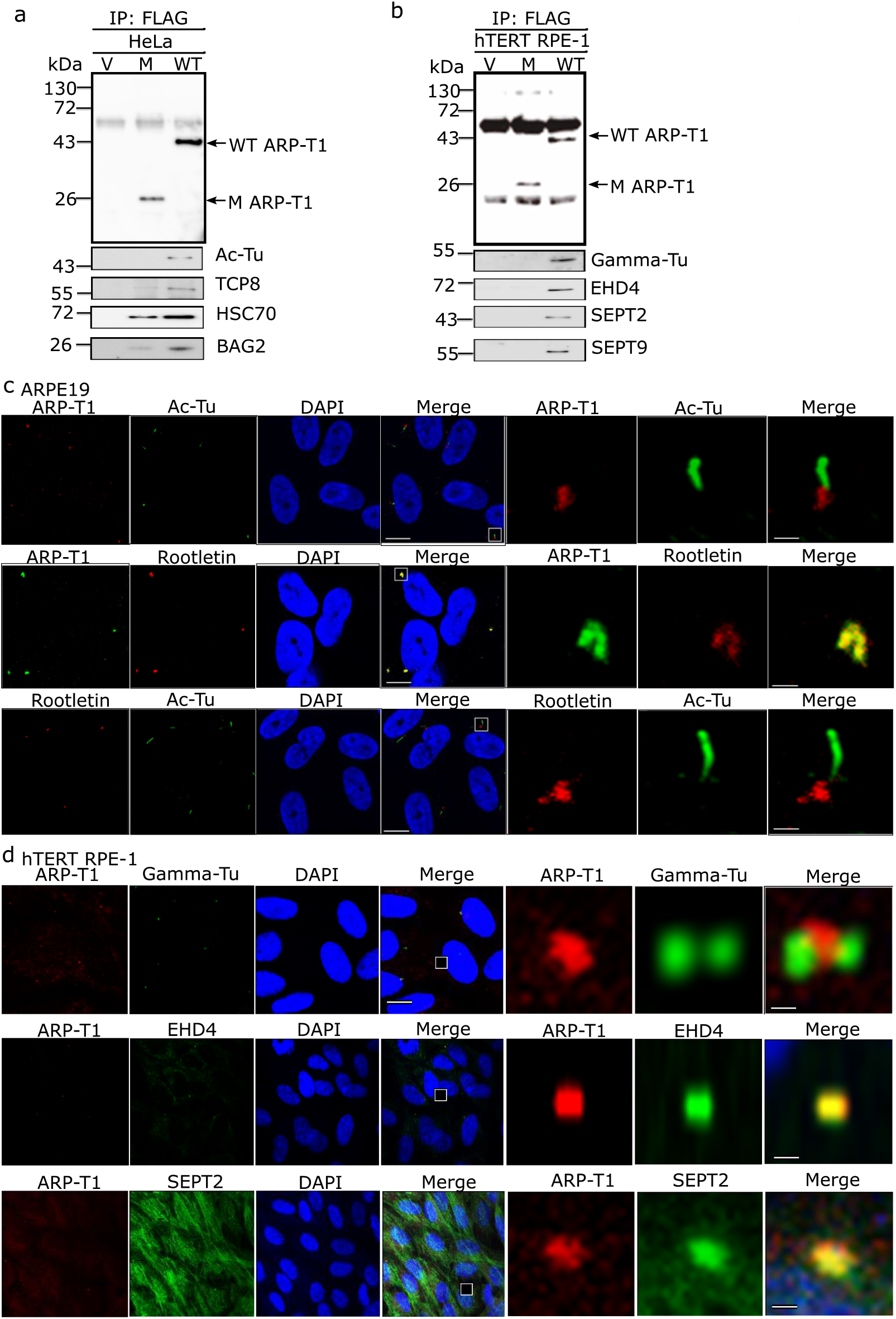
ARP-T1 interacts with proteins involved in ciliary machinery. **a**,**b**, HeLa (**a**) and hTERT-RPE1 (**b**) cells were transduced with lentiviral vectors, empty vector (V), *ACTRT1* mutant (M) and *ACTRT1* WT (WT), and immunoprecipitated (IP) with anti-FLAG monoclonal antibody M2-conjugated agarose, and analyzed by immuno-blot with indicated antisera. **c**, Immunofluorescence stainings of ARP-T1, acetylated-tubulin and rootletin in 35 days of serum-starved ARPE19 cells. Nuclei are stained with DAPI. Scale bar, 5 μm. Higher magnifications of the boxed area are shown on right three panels. Scale bar, 1 μm. **d**, Immunofluorescence staining of ARP-T1, gamma-tubulin, EHD4, and septin 2 in 48 h of serum-starved hTERT-RPE1 cells. Nuclei are stained with DAPI. Scale bar, 5 μm. Higher magnifications of the boxed area are shown on the right three panels. Scale bar, 1 μm.

We analyzed ciliogenesis, which was induced by serum starvation and differentiation, in ARPE-19 and hTERT-RPE1 cells. ARP-T1 localizes to the ciliary base when we co-stained ARP-T1 with acetylated-tubulin, which stains ciliary axoneme. ARP-T1 co-localized with rootletin, a major basal body protein, in PC (Fig. 3c and Extended data Fig. 1d in HaCaT cells). ARP-T1 interacts and forms a complex with gamma-tubulin, which localizes in the vicinity of the basal body (Fig. 3d and Extended data Fig. 1e in HaCaT cells). We also confirmed that ARP-T1 co-localizes with EHD4 and septin 2 (Fig. 3d). We used proximity ligation assays to further confirm these interactions (Extended data Fig. 1f and 1g). To reinforce these results, we chose to overexpress EHD4 and then locate ARP-T1. We transfected hTERT-RPE1 cells with EHD4 (EHD4) and vector control (V) tagged with V5. After immunoprecipitation using anti-V5 beads and immunoblot, we detected ARP-T1 in the EHD4 samples but not in the vector control (Extended data Fig. 1h). In the same cells, we confirmed that ARP-T1 co-localizes with EHD4 (Extended data Fig. 1i).

### ARP-T1 associates with a ciliopathy in BDCS

Based on the localization of ARP-T1 to the basal body and the implication of septin 2 in ciliary length control ^24^, we analyzed the ciliary structures in the samples from BDCS patients (*ACTRT1* 547_548insA, mutations B2, mutation A3, mutation CNE12)^6^ comparing to normal hair follicles and samples from a sporadic BCC patient. First, we observed the ciliary structures by staining with rootletin and acetylated-tubulin (Fig. 4a, top) and confirmed the co-localization of ARP-T1 and rootletin (Fig. 4a bottom, individual stainings with rootletin, acetylated-tubulin, or ARP-T1 are shown in Extended data Fig. 2 and 3). Strikingly, the ciliary length (Fig. 4b) and fluorescence intensity of rootletin and ARP-T1 (Fig. 4c,d) were significantly reduced in the BDCS samples compared to normal hair follicles and sporadic BCC. Ciliary length, rootletin fluorescence and ARP-T1 fluorescence were particularly reduced in *ACTRT1* 547_548insA (insA in Fig. 4b-f) and B2 mutation while less prominently reduced in mutations A2 and CNE12. We also found significant correlations between ARP-T1 fluorescence and ciliary length (Fig. 4e) and between ARP-T1 fluorescence and rootletin fluorescence (Fig. 4f). We concluded that BDCS, inherited BCC, unlike sporadic BCC, is a novel ciliopathy and the role of ARP-T1 as a part of cilia is to prevent the disease.

**Figure 4.**
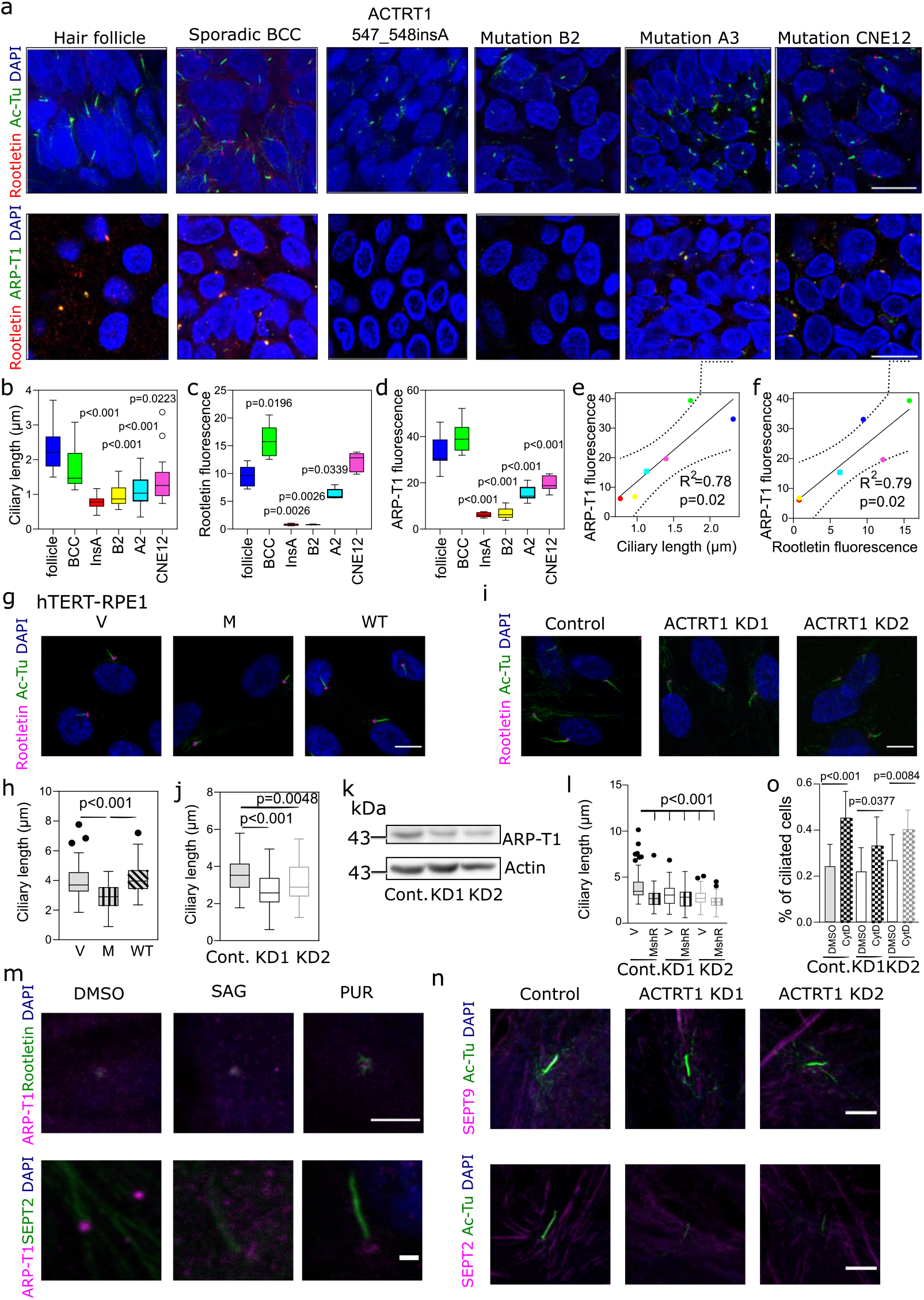
The Bazex-Dupré-Christol syndrome is a ciliopathy caused by ARP-T1 <u>loss of function,</u> and knock-down of *ACTRT1* in hTERT-RPE1 cells induces resorption of primary cilia. **a**, Representative immunofluorescence images using acetylated-tubulin (green) and rootletin (red) (left), and ARP-T1 (green) and rootletin (red) (right) in hair follicle, sporadic BCC and 4 BDCS (*ACTRT1* 547_548insA, mutation B2, mutation A3, mutation CNE12 ^6^). Cell nuclei are stained with DAPI (blue). Scale bar, 5 μm. **b**, Quantification of the ciliary length from 3D confocal immunofluorescence microscopy images. **c**,**d**, Quantification of the relative fluorescence intensity of rootletin (**c**, N=5) and ARP-T1 (**d**, N=10) on the ciliary rootlet. **e**,**f**, Correlation between the ARP-T1 fluorescence and ciliary length (**e**), and between the ARP-T1 and rootletin fluorescence (**f**). <u>**g**, Immunofluorescence stainings of acetylated-tubulin (green) and rootletin (pink) in 48 h serum starved hTERT-RPE1 cells. expressing an empty vector (V), or *ACTRT1* mutant (M), or *ACTRT1* WT (WT). Cell nuclei are stained with DAPI (blue). Scale bar, 10 μm. **h**, Quantification of ciliary length of **g**. **i**, Immunofluorescence stainings of acetylated-tubulin (green) and rootletin (pink) in 48 h serum-starved control and *ACTRT1* KD hTERT-RPE1 cells. Cell nuclei are stained with DAPI (blue). Scale bar, 10 μm. **j**,**k**, Quantification of the ciliary length (**j**) and ARP-T1 protein level (**k**, N=3) in control (Cont.) and *ACTRT1* KD hTERT-RPE1 cells. **l**, Quantification of ciliary length in 48 h serum-starved control and *ACTRT1* KD hTERT-RPE1 cells expressing an empty vector (V), or *ACTRT1* mutant resistant to shRNA (MshR). **b**-**d,h,j,l**, Results are presented as Tukey box-plot. Black circles represent outliers. **m**, Representative immunofluorescence stainings of ARP-T1 (pink) and rootletin (green, top) or septin 2 (green, bottom) upon treatment with SAG or purmorphamine (PUR) in hTERT-RPE1 cells under differentiating condition. Scale bar, 5 μm (top) or 1 μm (bottom). **n**, Immunofluorescence stainings of acetylated-tubulin (green) and septin 9 (pink, top) or septin 2 (pink, bottom) in 48 h serum-starved control and *ACTRT1* KD hTERT-RPE1 cells. Scale bar, 1 μm. **o**, Percentage of ciliated control and *ACTRT1* KD hTERT-RPE1 cells under proliferative condition, after treatment with cytochalasin D (CytD). Data are presented as means of the percentage +/−SD.</u>

<u>To study the role of ARP-T1 in controlling ciliary length *in vitro*, we overexpressed *ACTRT1* WT and *ACTRT1* 547_548insA in hTERT-RPE1 cells. The length of PC was quantified based on acetylated-tubulin staining. ARP-T1 mutant overexpression reduced the ciliary length, approximately by 25% (Fig. 4g and quantified in Fig. 4h, individual stainings are in Extended data Fig. 4a). We then investigated more closely the role of ARP-T1 with *ACTRT1* silencing in hTERT-RPE1 cells. Two different *ACTRT1* shRNAs efficiently knocked down the expression of *ACTRT1* at mRNA level (65% for KD1 and 50% for KD2; shown in Extended data Fig. 4c) and at ARP-T1 level (30%, Fig. 4k and Extended data Fig. 4d). PC in *ACTRT1*-depleted cells were shorter than in control cells (25% reduction; Fig. 1i,j). These results were confirmed in HaCaT cells, in which ciliary length was halved with *ACTRT1* KD1 and KD3 (Extended data Fig. 5a-c). To further confirm the role of ARP-T1 mutant in the control of ciliary length, we expressed *ACTRT1* 547_548insA resistant shRNA (MshR) in control and *ACTRT1* KD cells. ARP-T1 MshR expression leads to an identical reduction of ciliary length in control and KD cells (shown by the quantification in Fig. 4l, stainings are shown in Extended data Fig. 6a). These results confirmed that ARP-T1 loss of function and *ACTRT1* silencing both lead to the reduction of the ciliary length.</u> Therefore, ARP-T1 is required to maintain the normal length of cilia *in vitro*. In summary, ARP-T1 is located in ciliary basal body and regulates ciliogenesis both *in vivo* and *in vitro.* Absence of ARP-T1 gives rise to an epidermal ciliopathy.

**Figure 5.**
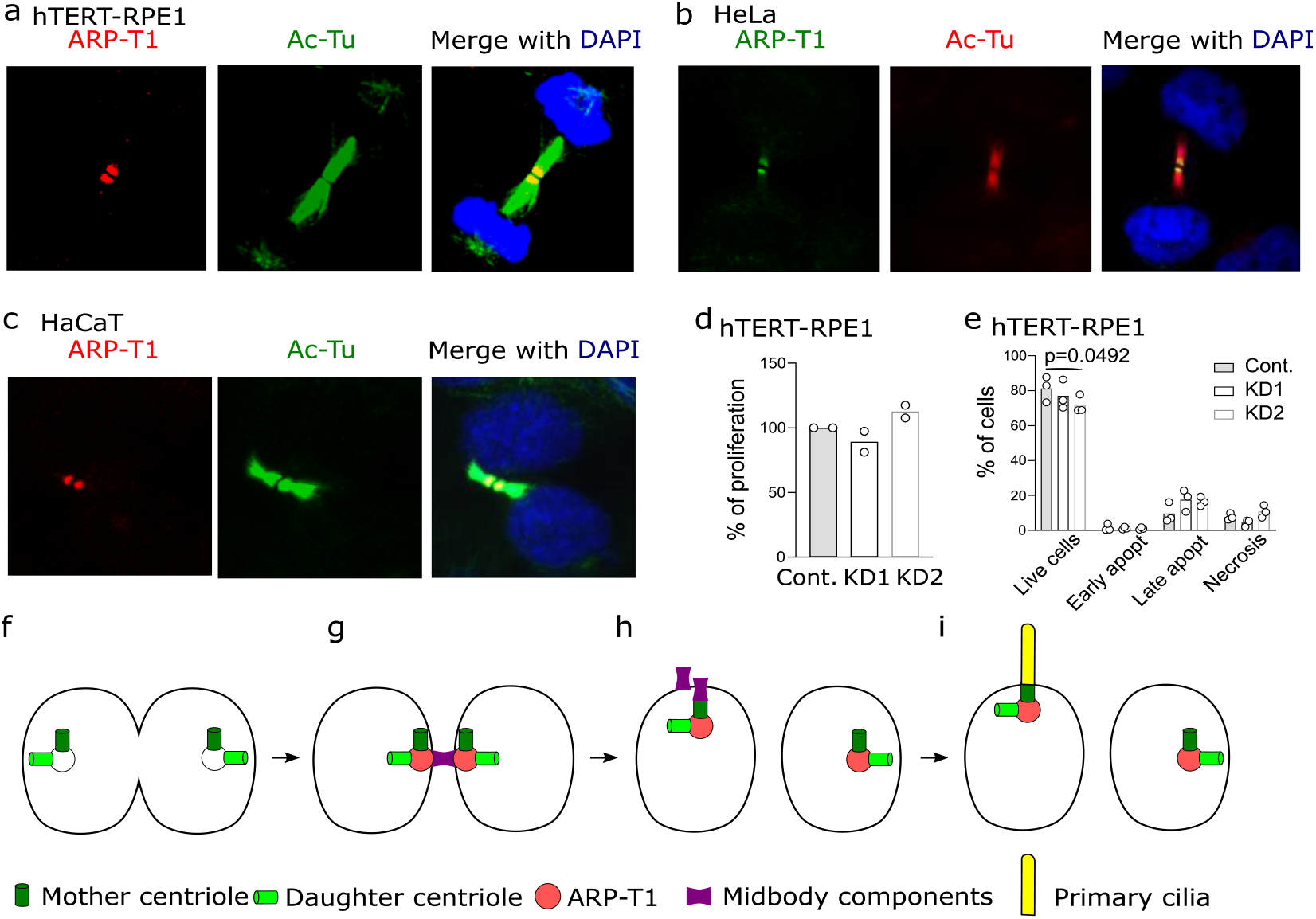
ARP-T1 localizes to midbody during cytokinesis. **a**, Immunofluorescence stainings of ARP-T1 (red) and acetylated-tubulin (green) in hTERT-RPE1 cells. Cell nuclei are stained with DAPI (blue). **b**, Immunofluorescence stainings of ARP-T1 (green) and acetylated-tubulin (red) in HeLa cells. Cell nuclei are stained with DAPI (blue). **c**, Immunofluorescence stainings of ARP-T1 (red) and acetylated-tubulin (green) in HaCaT cells. Cell nuclei are stained with DAPI (blue). <u>**d**,**e**, Proliferation (**d**) and apoptosis (**e**) analyses of control (Cont.) and *ACTRT1* KD hTERT-RPE1 cells. Data are presented as means of the percentage. Each open circle represents one independent experiment. **f**-**i**</u>, Model for ARP-T1 travelling from midbody to the PC.

### ARP-T1 favors for septin 2 localization in the axoneme, without affecting the actin cellular network

<u>PC is essential for HH pathway, and we described that ARP-T1 is localized at its basis and is necessary for the full length of the axoneme. Thus, we investigated the localization of ARP-T1 and its interacting proteins during HH activation. We found that ARP-T1 remains at the cilia basal body upon SMO activation, with SAG and purmorphamine, according to the co-localizations with rootletin and septin 2 (Fig. 4m). We also showed that remaining ARP-T1, in the *ACTRT1* KD cells, still co-localizes with the rootletin, and septins 2 and 9 (Extended data Fig. 6b) and that septin 9 localized to the axoneme, septin filaments and the base of the PC in control and *ACTRT1* KD cells under differentiation (Fig. 4n top). Therefore, ARP-T1 decrease does not affect neither its interactions with rootletin and septins nor septin 9 localization. Unexpectedly, we observed that septin 2, normally localized in the axoneme under serum-starvation, is no longer there in *ACTRT1* KD cells (Fig. 4n bottom). We were able to detect septin 2 in its other locations, i.e. septin filaments and the base of the PC.</u>

<u>Finally, as the actin cytoskeleton network is involved in ciliogenesis and as we found interactions between ARP-T1 and several proteins associated with this network (cortactin, septins, ARP2), we studied the actin cytoskeleton in *ACTRT1* KD cells and its involvement in cilia formation. We did not find any difference in actin filaments comparing *ACTRT1* KD and control cells (unpublished data). Moreover, cytochalasin D-induced ciliogenesis ^25^ was not blocked by silencing of *ACTRT1* (Fig. 4o), indicating that the actin cytoskeletal network is not deficient in these cells and is unlikely to be the cause of ciliary shortening.</u>

### ARP-T1 is located in the midbody in dividing cells

It was reported that septin 2 acts as a scaffold for myosin II and its kinases at the cleavage furrow ^26^ and that septin 9 mediates midbody abscission during cytokinesis ^27^. Therefore, we searched for the localization of ARP-T1 in dividing cells. ARP-T1 localized at the midbody in hTERT-RPE1 cells (Fig. 5a), HeLa cells (Fig. 5b) and HaCaT cells (Fig. 5c). <u>To determine whether ARP-T1 was implicated in cell division, we performed proliferation and apoptosis assays. Using EdU, an analog of thymidine, we found that ARP-T1 is not necessary for proliferation because control and *ACTRT1* KD cells incorporated identical amounts of EdU (Fig. 5d). We then analyzed apoptosis of these cells. ARP-T1 also has no effect on the different stages of apoptosis, although we observed a slight decrease in the live cell population with KD2 (Fig. 5e). In our conditions, *ACTRT1* silencing has no obvious impact on cell division.</u> Postmitotic midbody is directly implicated in primary ciliogenesis. After cytokinesis, the midbody remnant moves along the apical surface, proximal to the centrosome at the center of the apical surface where the PC emerges. The midbody remnant carries RAB8, IFT20, IFT88 and exocyst subunits required for ciliogenesis. If the remnant is removed, primary ciliogenesis is greatly impaired ^28^. Midbody remnant licenses PC formation ^29^. Both midbodies and PC contain acetylated tubulin, many proteins in the midbody can also be found at the base of the cilium in the centrioles ^30^ (Fig. 5f-i).

## DISCUSSION

Here, we report that ARP-T1 is a <u>ciliogenesis</u> protein located at the basal body of PC. BDCS tumors and *ACTRT1*-deficient cells show shortened PC. Thus, BDCS is a novel and first ciliopathy implicated in skin cancer caused by mutations of *ACTRT1* <u>or its enhancer RNA elements.</u> ARP-T1 supports intact cilia and <u>controls</u> proper ciliogenesis<u>, potentially through septin 2 involvement</u>. ARP-T1 is highly expressed during epidermal differentiation, is stabilized by protein kinase C and *ACTRT1* expression is regulated by PKC delta and by a non-canonical HH pathway.

*ACTRT1*/ARP-T1 regulation has not been described yet. Here, we show that HH and mTORC2/PKC delta pathways positively regulate *ACTRT1* at the mRNA level. HH signaling is activated upon cell differentiation, at the cilium, and is known to induce canonical and non-canonical signaling pathways. As ARP-T1 was previously reported to activate the HH signaling ^6^, it was important to decipher a possible feedback loop with this pathway. We found that SMO activation favors *ACTRT1* mRNA <u>and ARP-T1 protein</u> expression. However, a blockade with vismodegib, used in the clinic to treat BCC patients, could not reverse purmorphamine-induced *ACTRT1* mRNA expression. We concluded that *ACTRT1* <u>mRNA</u> is regulated by a non-canonical HH signaling. We next unraveled which phosphorylation pathway was implicated in this non-canonical HH signaling. MAP kinase inhibition with PD98059 had no effect on *ACTRT1* <u>mRNA</u> expression. By using PI3K/mTORC1/S6K and mTORC2/PKC delta inhibitors, we found that the proliferating signaling PI3K/mTORC1/S6K limits *ACTRT1* <u>mRNA</u> expression while its differentiating counterpart mTORC2/PKC delta ^28^ favors it. These findings highlight the role of HH and mTORC2/PKC delta pathways in *ACTRT1*/ARP-T1 regulation.

By MS analysis, we identified a machinery of molecular ARP-T1 interactors involved in PC formation (see Table 1): (1.) Chaperon containing TCP-1 (CCT/TRiC), consists of two identical stacked rings, each containing eight subunits (TCP1-8). This complex folds various proteins including actin and tubulin. It was shown that CCT/TRiC family chaperonin localizes to centrosomes, forms a complex with BBS proteins BBS6, BBS10 and BBS12, and mediates the assembly of the BBSome, involved in ciliogenesis regulating transports of vesicles to the cilia ^29^. This BBSome protein complex binds to Rabin 8, the GTP/GDP exchange factor, for the small GTPase RAB8. RAB8 localizes to the cilium, is required for ciliogenesis, and mediates the docking and fusion of vesicles. It was proposed that the BBSome acts upstream of RAB8 in this vesicular transport to the cilium ^30^. BBS causing IFT27 mutations point to a link of the BBSome with IFT where this small GTPase might be involved in delivering cargo from the BBSome to the IFT ^31^ (2.) CCT and HSC70 form a stable complex and this interaction was suggested to be used to deliver the unfolded substrate from HSC70 to the substrate-binding region of CCT ^32^. HSC70 interacts with IFT88 ^33^. Bcl2-associated athanogen (BAG2) protein forms a chaperone complex with HSC70 and regulates their protein folding activity ^34^; (3.) The Eps15 homology domain (EHD) family of proteins, composed of EHD1-4, is associated with RAB8 and RAB11 membranes and regulates endosomal membrane trafficking ^35^. EHD4 locates to the transition zone of cilia and plays a role in protein transport to cilia and ciliogenesis. *Ehd4* deficient mice show a strong phenotype in skin, kidney and testis with small testis, reduced sperm motility, sperm number and germ cells, such that *Ehd4*−− mice are subfertile ^36^. *Ehd4* deficient mice suffer from hypotrichosis at the dorsal back and have a PC phenotype (Dr. Markus Plomann, unpublished data); (4.) Septin 2 localizes to the base of the axoneme. Cells completely depleted of septin 2 lack a cilium whereas cells partially depleted at the base of PC have a significantly shortened cilium than control ^24^. Septin 2 interacts with HSC70 ^37^. Septin 9 has been shown to interact with microtubules ^38^ and localizes to the ciliary pocket region ^39^. PC in septin 9 depleted cells are shorter than that in control cells ^39^. Gamma-tubulin associates with centrosomes and localizes to the vicinity of basal body ^40^.

Overall, our MS analysis results point to a function of ARP-T1 in the ciliary basal body at the crossroads of vesicle trafficking, protein folding and the cytoskeleton (Supplementary Table 1). The ciliary pocket or periciliary PM serves as an interface with the actin cytoskeleton, which is important for the movement of the basal body to the PM and for vesicular and endosomal trafficking ^10,41,42,43^. We hypothesize that ARP-T1 is functional at this interface and may stabilize the cytoskeletal matrix.

Functional genomic screening showed that proteins involved in actin dynamics and endocytosis are important for ciliogenesis. Depletion of gelsolin family proteins (GSN and AVIL), which regulate cytoskeletal actin organization by severing actin filaments, reduced the number of ciliated cells and silencing of *ACTR3* (ARP-3), a major constituent of the ARP2/3 complex involved in nucleating actin polymerization, increased cilium length ^44^. On the other hand, it was reported that the actin cytoskeleton is disrupted in KD cell lines targeting ciliary proteins. For example, KD cells of *Ahi1*, whose human ortholog is mutated in Joubert syndrome, showed disorganizing and decrease in actin filaments ^45^. ARP-T1 has 49 % identity with β-actin and 40%, 37%, 29% identity with ARP-1, ARP-2, ARP-3, respectively. ARP-T1 as a <u>basal body</u> protein is involved in a positive regulation of ciliogenesis. Similar to ARP-T1, some cancer-associated mutations result in ciliary resorption: (1.) Von Hippel-Lindau (VHL) disease is characterized by the development of premalignant renal cysts, and arises because of functional inactivation of VHL tumor suppressor protein (pVHL). pVHL maintains the structural integrity of the PC for suppression of uncontrolled proliferation of kidney epithelial cells and cyst formation ^46^. (2.) It was shown by electron microscopy that glioblastoma cells and tumors have immature PC and basal body/centrioles ^47,48^. (3.) Mutations in Budding uninhibited by benzimidazole-related 1 (BUBR1), a molecule of spindle assembly checkpoint, cause premature chromatid separation (mosaic variegated aneuploidy), cancer disposition and impaired ciliogenesis ^49^. <u>Despite all these clues, we could not detect a difference in the actin filaments between *ACTRT1* KD and control cells, and our cytochalasin D-induced ciliogenesis experiments did not allow us to confirm a role of the actin cytoskeletal network in the shortened cilia observed in silenced *ACTRT1* cells.</u>

<u>Diffusion barrier is another cellular structure essential for the integrity of the PC ^38^. Septins are key components of the diffusion barrier at the base of the PC, forming ring-like structures and being transported to the axoneme. The involvement of septin 2 in ciliogenesis was particularly studied these last years ^24,39,50^. In an attempt to understand how the knockdown of *ACTRT1* led to a shortening of ciliary length, we observed that septin 2 was absent from the axoneme in *ACTRT1* silenced cells, but preserved in septin filaments and in the base of the PC. Such a situation was different for septin 9, whose location remained unchanged, i.e. in the axoneme, septin filaments and the base of the PC. These results suggest that the ciliogenesis defect observed in *ACTRT1* deficient cells could be due to a default in the diffusion barrier mediated by septin 2. We are currently investigating the link between septin 2 and ARP-T1 in cilia formation, notably using SMO in/out assay ^51^ and FRAP experiments ^39^. These results will be part of a forthcoming publication.</u>

<u>Septins are not only essential for the diffusion barrier at the base of PC, but also participate in cell division, where they are located in the midbody ^27^. Our investigations have shown that ARP-T1 is also found in the midbody of dividing cells, and that it could therefore have an impact on cell division. Nevertheless, cell proliferation and apoptosis were not affected by the decrease of ARP-T1, and thus ARP-T1 at the midbody level has no effect on cell division. This was not unexpected, as Pejskova et al. ^52^ recently described that KIF14, a protein involved in ciliogenesis, was also localized in the midbody without any impact on cell division.</u>

With our results, we propose a model where ARP-T1 leads primary ciliogenesis via direct interaction with proteins in ciliary machinery. First, ARP-T1 is localized in the mitotic spindle through the interaction with gamma-tubulin, which is present in between the mother centriole and the daughter centriole (Fig. 5f). As the cytokinesis proceeds, the complex of ARP-T1 and two centrioles moves to the central spindle where the midbody is formed by recruiting septins (Fig. 5g). The midbody cleaves one side of the intracellular bridge and remains tethered to one of the daughter cells. The complex of ARP-T1, two centrioles and midbody move towards the apical surface of daughter cells where PC will be formed (Fig. 5h,i).

Dynamic HH signaling has been identified in mammalian post-natal testis, and HH pathway components PTCH1, SUFU and GLI1 were identified in spermatogonia, spermatocytes and spermatids ^53^, where spermatids express high levels of SUFU to repress HH activity ^54^. It was previously reported that ARP-T1 binds to the *GLI1* promoter and inhibits *GLI1* mRNA expression signaling ^6^. In the current study, we showed that the elevated level of <u>*ACTRT1* mRNA</u> with purmorphamine was not decreased by the treatment with purmorphamine and vismodegib (Fig. 2c-e). <u>Nevertheless, this regulation is absent at the ARP-T1 protein level, as purmorphamine-induced ARP-T1 expression was counteracted by the addition of vismodegib (Fig. 2c).</u> Thus, <u>we hypothesize that</u> *ACTRT1* functions in a SMO-independent HH pathway <u>and that post-transcriptional modifications, as shown for pKC, regulate ARP-T1 function.</u> The activation of a non-canonical HH pathway may explain the apparent paradox that we observed shortened PC both in BCC from BDCS and in *ACTRT1* KD cells. While loss of PC is observed in diverse cancers where it correlates with worse prognosis, their retention is paramount for medulloblastoma and BCC development ^9^. Nevertheless, SMO-independent operation occurs both in medulloblastoma ^12^ and BCC mouse models ^13^, and was recently linked to development of resistance to SMO-inhibition ^55^. This suggest that BDCS is a model for the study of vismodegib resistance in BCC and medulloblastoma.

<u>Functions of the PC, including activation of the HH signaling, are dependent on the ciliary length and on proteins involved in ciliogenesis, such as proteins located to the basal body ^56^. In addition to this, ARP-T1 was reported to regulate the HH pathway ^6^. Here, as we demonstrated that *ACTRT1* silencing is associated with a shortened PC, we assessed the expression of HH proteins in our deficient cells. *GLI1* and *PTCH1* mRNA expression tends to increase but not significantly in *ACTRT1* KD cells (Extended data Fig. 6c,d), suggesting that our silencing might not be sufficient to correctly activate the HH signaling, or that another hit might be needed for a proper activation. In order to look for a second hit necessary for a strong activation of *GLI1*, we studied in depth the transcriptomic analysis performed in Bal et al. ^6^, and we unraveled a higher expression of *ARHGAP36* mRNA in BDCS samples, with an approximately 17-fold increase. *ARHGAP36* gene is located to the X chromosome, in the close proximity of *ACTRT1* gene, and is known to activate the HH pathway in medulloblastoma ^57^. ARHGAP36 might contribute to *GLI1* activation in BDCS patients. Another gene of interest is *SMARCA1*. Indeed, *SMARCA1* expression is often lost in gastric cancer cells due to methylation ^58^ and in soft tissue sarcomas ^59^. This gene is located next to *ACTRT1* gene, and it cannot be excluded that mutations in the RNA elements regulating *ACTRT1* do not also affect the expression of *SMARCA1*, which expression is 1.8-fold higher in BDCS patients ^6^. *SMARCA1* encodes for the probable global transcription activator SNF2L1, which act as a chromatin remodeler and interacts with actin. Loss of expression of *SMARCA1* increased proliferation of gastric cancer cells, giving it a tumor suppressor role ^58^. A whole exome sequencing of BDCS tumors is presently on going and we are currently studying the link between ARHGAP36, SNF2L1, and ARP-T1. These results will be the focus of a future publication.</u>

Our results show that ARP-T1 acts as a direct or indirect actor in a non-canonical HH pathway and bridges between actin cytoskeleton organization involved in vesicle transport at the centrosome, the centrioles and the basal body, and the formation of PC to prevent the pathogenesis of BDCS. Above all, our studies shed light on how ciliogenesis is controlled in carcinogenesis by ARP-T1. ARP-T1 could be a new target for novel therapeutic approaches in BDCS and BCC, the most frequent human cancer. This appears particularly important since loss of PC is a recognized mechanism of resistance to SMO inhibitors in medulloblastoma and BCC ^55,60^. <u>Finally, Glaessl et al. ^61^ described BDCS BCCs to be more aggressive and prone to relapse. As BDCS patients have a shortened PC, we may hypothesize that BDCS BCCs are a more advanced form of BCC.</u>

## Online METHODS

### Cell Culture and samples

Normal human epidermal keratinocytes (NHEK, established from neonatal foreskin in our laboratory) and immortalized keratinocytes, HaCaT cells (Invitrogen, Basel, Switzerland), were grown in EpiLife medium (Gibco, Invitrogen) with Human Keratinocyte Growth Supplement (Gibco, Invitrogen), 10 μg/ml gentamicin and 0.25 μg/ml amphotericin B (Gentamicin / Amphotericin B solution, Gibco, Invitrogen). Spontaneously immortalized adult retinal epithelial cell line 19 (ARPE-19) and immortalized hTERT-RPE1 cells (ATCC, Manassas, VA) were cultured in Dulbecco’s Modified Eagle’s Medium (DMEM) High Glucose, GlutaMAX™, Pyruvate (Gibco, Invitrogen) or DMEM/F12 (ATCC) with 0.01 mg/ml hygromycin B (Sigma-Aldrich, St Louis, MO), both media supplemented with 10% fetal bovine serum (FBS), 100 U/ml penicillin and 100 μg/ml streptomycin (BioConcept AG, Allschwil, Switzerland). HeLa and HEK293T (Invitrogen) were cultured in DMEM supplemented with 10% FBS, 100 U/ml penicillin and 100 μg/ml streptomycin. The cells were incubated at 37°C in a 5% CO_2_ atmosphere.

To induce cilia formation, NHEK and HaCaT were cultured up to 7 days in EpiLife media supplemented with 2 mM CaCl_2_. hTERT-RPE1 and ARPE-19 cells were cultured 48 h or 35 days respectively in media containing 0.2% FBS.

Bazex-Dupré-Christol syndrome samples were collected and prepared as previously described ^6^.

### Lentivirus <u>production and transduction</u>

<u>Lentivirus particles (*LVs*)</u> for *ACTRT1* shRNA were produced by Sigma-Aldrich, in pLKO backbone vector. ShRNA sequences are for *ACTRT1* KD1: 5’ CCG GGC CTG GTT TCT ACC TGT CTA ACT CGA GTT AGA CAG GTA GAA ACC AGG CTT TTT G 3’, and *ACTRT1* KD2: 5’ CCG GGT GCC TTT AGC AAG ACT TAA TCT CGA GAT TAA GTC TTG CTA AAG GCA CTT TTT TG 3’; <u>*ACTRT1* KD3: CCG GCA TGA CCT CTA TGA GCA GTT TCT CGA GAA ACT GCT CAT AGA GGT CAT GTT TTT G.</u> Control is an empty vector with same selection.

<u>LVs for *ACTRT1* overexpression were produced with calcium phosphate transfection method: *ACTRT1* wild-type (*WT*) and mutant containing the 547-548InsA mutation constructs were produced as previously described ^6^. HEK293T cells were transiently co-transfected with psPAX2 (Addgene, Cambridge, MA) and pMD2.G (Addgene) and empty vector or *ACTRT1* WT or mutant to produce LVs. LVs were harvested 48 h later and the titer was determined in HeLa cells ^62^. *ACTRT1* 547-548InsA construct was mutated using QuikChange Multi Site-Directed Mutagenesis Kit (200514, Agilent Technologies, Basel, Switzerland) in the KD1 target (T599G, C602T, C605T) and reverse KD2 target (A341G, A344C, T556G) with following primers (Microsynth, Balgach, Switzerland): KD1 target GCCTGGGTTTTATCTGTCTAA, KD2 target GATTCAGTCTGGCCAAAGGCAC.</u>

HeLa, hTERT-RPE1 and HaCaT cells were transduced using polybrene (8 ug/mL, TR1003, Millipore), with LVs O/N at 37°C. The next day, LVs were removed and cells grew in their medium. <u>When necessary, s</u>election with puromycin (2.5 ug/mL, 540411, Calbiochem) or blasticidin (7.5 ug/mL, 15205, Sigma-Aldrich) started 24 h later.

### Drug Treatments

Cells were treated 24 h with SMO agonists 200 and 400 nM SAG (566660, Calbiochem) and 3 μM purmorphamine (4551, Tocris bioscience), or antagonist 5 μM vismodegib (S1082, Selleckchem), or kinase inhibitors: 10 μM GF109203X (ALX-270-049, Enzo Life Sciences), 10 μM SB431542 (1614, Tocris bioscience), 20 μM PD98059 (9900, Cell Signaling Technologies), 5 μM SP600125 (270-339-M005, Alexis Biochemicals), 10 μM SB216763 (1616, Tocris bioscience), 200 nM Rapamycin (R0161, LKT Laboratories), 50 μM LY294002 (70920, Cayman Chemical), 5 μM AG1478 (270-036-M001, Alexis Biochemicals), 3 μM Gö6976 (12060S, Cell Signaling Technologies), 10 μM Rottlerin (1610, Tocris bioscience), 20 μM ZIP (2549, Tocris bioscience, kind gift from Prof. Anthony Oro, Stanford University School of Medicine).

<u>To induce cilia in proliferating condition, hTERT-RPE1 cells were treated for 16 h with 50 nM cytochalasin D (250255, MERCK).</u>

### Western blot

NHEK, HaCaT, ARPE-19, and hTERT-RPE1 cells were harvested in ice-cold FLAG (50 mM Tris/HCl pH 7.5, 150 mM NaCl, 1 mM EDTA, 1% Triton X-100, and protease inhibitor cocktail [Complete MiniTM tablette, Roche Diagnostics]) or RIPA (50 mM Tris/HCl pH 7.4, 150 mM NaCl, 12 mM Na deoxycholate, 1% NP-40, 0.1% SDS, protease inhibitor cocktail, and phosphatase inhibitor cocktail [PhosSTOP, PHOSS-RO, Roche]) lysis buffers for 30 min on ice with extensively pipetting every 10 min, then spun for 10 to 20 min at 12,000 x *g* at 4°C. Supernatants were collected, heated for 5 min at 85°C and centrifuged. Total protein concentration was determined by BCA assay (PierceTM BCA Protein Assay Kit; Thermo Scientific, Waltham, MA, USA). Fifteen ug of proteins were loaded onto a 10% SDS-PAGE gel and electroblotted onto a nitrocelulose or PVDF membrane (Hybond ECL; Amersham, UK). Membranes were incubated overnight at 4°C with the following antibodies: anti-ARP-T1 (1:1000, SAB1408334, Sigma-Aldrich / 1:2000, GP-SH6, Progen), anti-keratin 10 (1:200, MS-611-P0, Thermo Scientific), anti-IFT88 (1:1000, 13967-1-AP, Proteintech, UK), anti-actin (loading control, 1:5000, A2066, Sigma-Aldrich) and anti-alpha-tubulin (loading control, 1:5000, T9026, Sigma-Aldrich). After washes in TBS-T, 1 h incubation at room temperature (RT) with HRP-secondary antibodies (anti-mouse [1:5000, NA931V, GE Healthcare UK Limited], anti-rabbit [1:5000, NA934V, GE Healthcare], anti-guinea pig [1:5000, ab97155, Abcam]), and revelation with chemiluminescence (ECL Prime, Amersham / WesternBright Quantum, Advansta, Witec, Sursee, Switzerland), images were acquired using the Luminescent Image Analyzer LAS-4000 mini (Fujifilm, Tokyo, Japan). Band intensity was analyzed with Image J software (National Institutes for Health, USA).

### Co-immunoprecipitation

Transduced HeLa, NHEK, and serum-starved hTERT-RPE1 cells were washed with PBS and harvested in the FLAG lysis buffer. Supernatants were incubated with 30 μL anti FLAG-beads (ANTI-FLAG^TM^ M2 Affinity Gel) overnight at 4°C and 1 h at RT on a rotator to pool-down ARP-T1-FLAG proteins. Beads were separated by centrifugation for 3 min at 5000 x *g*, washed five times with TBS, and eluted in 23 μL TBS and 7 μL 5x SDS-sample buffer at 85°C for 5 min. ARP-T1 precipitation was confirmed using ARP-T1 antiserum (1:2000, GP-SH6), and co-precipitated proteins were analyzed using anti-acetylated-tubulin (1:1000, T6793, Sigma-Aldrich), anti-TCP8 / TCP1 theta (1:500, PA5-30403, Thermo Scientific), anti-HSC70 (1:200, PA5-27337, Thermo Scientific), anti-BAG2 (1:100, PA5-30922, Thermo Scientific), anti-gamma-tubulin (1:500, ab11316, Abcam), anti-EDH4 (1:1000, kindly provided by Dr. Plomann<u>, unpublished antibody</u>), anti-septin 2 (1:2000, HPA018481, Sigma-Aldrich), anti-septin 9 (1:2000, HPA042564, Sigma-Aldrich).

### Proteomic analysis/ Mass Spectrometry

Transduced HeLa, NHEK, and serum-starved hTERT-RPE1 cells were washed with PBS and harvested in FLAG lysis buffer, and analyzed by the Proteomic Analysis Facility of University of Lausanne with the following protocol.

#### Gel separation and protein digestion

Protein samples were loaded on a 12% mini polyacrylamide gel and migrated about 2.5cm in non-reducing conditions. After Coomassie staining, gel lanes between 15-300 kDa were excised into 5-6 pieces, and digested with sequencing-grade trypsin (Promega) as described by Shevchenko and colleagues ^63^. Extracted tryptic peptides were dried and resuspended in 0.05% trifluoroacetic acid, 2% (v/v) acetonitrile for mass spectrometry analyses.

#### Mass spectrometry analyses

Tryptic peptide mixtures were injected on a Dionex RSLC 3000 nanoHPLC system (Dionex, Sunnyvale, CA, USA) interfaced via a nanospray source to a high resolution mass spectrometer based on Orbitrap technology: Orbitrap Fusion Tribrid or QExactive Plus instrument (Thermo Fisher, Bremen, Germany), depending on the experiments considered. Peptides were loaded onto a trapping microcolumn Acclaim PepMap100 C18 (20mm x 100μm ID, 5μm, Dionex) before separation on a C18 reversed-phase analytical nanocolumn, using a gradient from 4 to 76% acetonitrile in 0.1% formic acid for peptide separation (total time: 65 min).

Q-Exactive Plus instrument was interfaced with an Easy Spray C18 PepMap column (25cm x 75μm ID, 2μm, 100Å, Dionex). Full MS survey scans were performed at 70,000 resolution. In data-dependent acquisition controlled by Xcalibur software (Thermo Fisher), the 10 most intense multiply charged precursor ions detected in the full MS survey scan were selected for higher energy collision-induced dissociation (HCD, normalized collision energy NCE=27%) and analysis in the orbitrap at 17’500 resolution. The window for precursor isolation was of 1.5 m/z units around the precursor and selected fragments were excluded for 60s from further analysis.

Orbitrap Fusion Tribrid instrument was interfaced with a reversed-phase C18 Nikkyo column (75μm ID x 15cm, 3.0μm, 120Å, Nikkyo Technos, Tokyo, Japan) or a custom packed column (75μm ID × 40cm, 1.8μm particles, Reprosil Pur, Dr. Maisch). Full survey scans were performed at a 120’000 resolution, and a top speed precursor selection strategy was applied to maximize acquisition of peptide tandem MS spectra with a maximum cycle time of 3 s. HCD fragmentation mode was used at a normalized collision energy of 32%, with a precursor isolation window of 1.6 m/z, and MS/MS spectra were acquired in the ion trap. Peptides selected for MS/MS were excluded from further fragmentation during 60 s.

#### Data analysis

MS data were analyzed using Mascot 2.5 (Matrix Science, London, UK) set up to search the Swiss-Prot (www.uniprot.org) database restricted to Homo sapiens taxonomy (UniProt, December 2015 version: 20’194 sequences). Trypsin (cleavage at K,R) was used as the enzyme definition, allowing 2 missed cleavages. Mascot was searched with a parent ion tolerance of 10 ppm and a fragment ion mass tolerance of 0.50 Da (Orbitrap Fusion) or 0.02 Da (QExactive Plus). Carbamidomethylation of cysteine was specified in Mascot as a fixed modification. N-terminal acetylation of protein and oxidation of methionine were specified as variable modifications.

Scaffold software (version 4.4, Proteome Software Inc., Portland, OR) was used to validate MS/MS based peptide and protein identifications, and to perform dataset alignment. Peptide identifications established at lower than 90.0% probability by the Scaffold Local FDR algorithm were filtered out. Protein identifications were accepted if they could be established at greater than 95.0% probability and contained at least two identified peptides. Protein probabilities were assigned by the Protein Prophet algorithm^64^. Proteins that contained similar peptides and could not be differentiated based on MS/MS analysis alone were grouped to satisfy the principles of parsimony. Proteins sharing significant peptide evidence were grouped into clusters.

The mass spectrometry proteomics data have been deposited to the ProteomeXchange Consortium ^65^ via the PRIDE ^66^ partner repository with the dataset identifier PXD016557 and 10.6019/PXD016557.

### Immunofluorescence

Cells grown on coverslips were fixed in 4% formaldehyde/PBS for 20 min at RT, then permeabilized in 0.02% Triton-X/PBS for 10 min. Coverslips were incubated with blocking buffer containing 1% FBS and 2% bovine serum albumin (*BSA*) for 1h. Primary antibodies (anti-ARP-T1 (1:100, SAB2103464 <u>or SAB1408334; or</u> 1:200, Progen GP-SH6), anti-acetylated-tubulin (1:1000, T6793), anti-rootletin (1:50, sc-67824, Santa-Cruz; 1:200, NBP1-80820, Novus), anti-gamma-tubulin (1:500, ab11316, Abcam), anti-EDH4 (1:200, kind gift from Dr. Markus Plomann, <u>unpublished antibody</u>), anti-septin 2 (1:200, HPA018481, Sigma-Aldrich), anti-septin 9 (1:200, HPA042564) were diluted in blocking buffer and applied on the cells for 2 h at RT or overnight at 4°C. Coverslips were next incubated with secondary antibodies (1:500, A11008, A11001, A21467, A11035, A11003, A11060, Invitrogen; 1:100, BA-9500, Vector Laboratories; 1:200, RPN1233V, GE) for 1 h at RT. Nucleus was stained with DAPI or Hoechst for 5 min at RT. Finally, coverslips were mounted onto slides with Dako mounting medium (S3023, Dako Schweitz AG, Baar, Switzerland) and examined with an inverted Zeiss LSM 700 laser scanning confocal microscope equipped with laser diode 405/488/555, SP490/SP555/LP560 emission filters, 2 PMT detectors and Zen2010 software (Zeiss, Feldbach, Switzerland).

Three-dimensional (3D) imaging technique was used to study ciliary imaging. Confocal images were captured at ~0.33-1μm interval using 63 x/1.40 oil objectives. Using Image J software, the 3D structure was deconvoluted from the Z-stack, and the length, intensity and prevalence of cilia were manually traced and measured. Both cell number and proportion of cells exhibiting PC were determined in approximately five representative fields (100 to 200μm^2^) for each experimental condition. The mean cilia prevalence was expressed as the percentage of ciliated cells. The average ciliary length was expressed in μm. The ciliary rootlet was selected using the freehand tool and the average area was expressed in μm^2^. The average pixel fluorescence intensity of proteins was quantified using the freehand tool to select the area of interest. Twenty to 50 cilia were measured for each experimental condition.

### RNA extraction and Real time quantitative PCR

Total RNA were harvested using the Qiagen RNeasy Mini Kit (Qiagen, Hombrechtikon, Switzerland) according to the manufacturer’s instructions. RNA quantity and quality were assessed using the NanoDrop ND-1000 Spectophotometer (Wilmington, USA). RNA were reverse-transcribed using the Primescript RT reagent kit (TakaRa, Saint-Germain-en-Laye, France) in 10 μL reaction on a T-Gradient Thermocycler (Biometra, Biolabo, Châtel-St-Denis, Switzerland). Quantitative PCR (*qPCR*) was performed on cDNA with Power SYBRGreen PCR Mastermix (Applied Biosystems, CA, USA) on an Applied Biosystems StepOne thermal cycler (Applied Biosystems, Life Technologies, CA, USA). Gene expression was assessed by using *ACTRT1* (QT00215642, Qiagen), *GLI1* (Fw 5’ AGA GGG TGC CAT GAA GCC AC 3’, Rev 5’ AAG GTC CCT CGT CCA AGC TG 3’, Microsynth), *PTCH1* (Fw 5’ GCT ACT TAC TCA TGC TCG CC 3’, Rev 5’ TCC GAT CAA TGA GCA CAG GC 3’, Microsynth) primers. *RPL13A* (Fw 5’GCA TCC CAC CGC CCT ACG AC 3’, Rev 5’ CTC TTT CCT CTT CTC CTC CA 3’, Microsynth) was used as endogenous control. Relative gene expression is normalized to the control and reported with a log2 scale.

#### Proliferation assay

<u>Proliferation was assessed using the Click-iT EdU Proliferation Assay for Microplates (C10499, Invitrogen). Briefly, cells were plated at 48% confluency in 96 well black microplate (CLS3603, Corning, MERCK), labelled with 10 μM EdU for 24h, then fixed and clicked according to the manufacturer’s instructions. Fluorescence was analyzed with a Mithras LB 940 reader (Berthold technologies, Bad Wildbad, Germany) and MikroWin 2010 software (Berthold technologies) using excitation filter 560 nm and emission filter 590 nm.</u>

#### Apoptosis assay

<u>Cells were harvested in Versene (15040, Gibco, ThermoFisher Scientific), then stained with APC Annexin V (550474, BD Pharmingen, San Jose, CA) and Annexin V Apoptosis Detection Kit I (556547, BD Pharmingen) according to the manufacturer’s instructions. Results were collected with a FACS Gallios I (Beckman Coulter, Nyon, Switzerland) and analyzed with FlowJo software (BD Life Sciences) at the Flow Cytometry Facility at Unil.</u>

### Statistics and Reproducibility

We used the one-way ANOVA or the Kruskal-Wallis test, depending on homogeneity of variances, to compare one independent variable in more than two groups. Linear regression was used to compare the correlation between two parameters.

To compare gene expression quantified by RT-qPCR, for each gene, all samples were normalized to the control, and log-2 transformed fold changes comparison to 0 were performed with a one-sample t test. To compare protein expression, all samples were normalized to the control, and fold changes comparison to 1 were performed with a one-sample t test.

<u>All experiments were performed at least three times.</u> All statistical analyses were performed using Prism GraphPad (v8, La Jolla, CA, USA).

## Data Availability

<u>Original immunoblots and immunofluorescence data are accessible on Zenodo repository with the dataset identifier 10.5281/zenodo.4317153 (all versions: 10.5281/zenodo.3666278).</u>

## SUPPLEMENTARY METHODS

### In situ proximity-mediated ligation assay (PLA)

<u>hTERT-RPE1 cells were seeded on coverslips, fixed and permeabilized. Duolink in situ PLA kit with anti-rabbit PLUS probe and anti-mouse or anti-goat MINUS probe (Sigma-Aldrich) was used according to manufacturer`s instructions: blocking for 30 min in a 37°C humidified chamber, incubation with primary antibodies (gamma-tubulin (1:500, ab11316), ARP-T1 (1:100, SAB2103464), rootletin (1:50, sc-67824), EHD4 (1:200, ab83859, Abcam), septin 2 (1:100, HPA018481), septin 9 (1:100, HPA042564) for 2 h at RT or overnight at 4°C, hybridization with PLA PLUS and MINUS probes (1:5 dilution) for 1 h at 37°C, ligation, amplification and final washes. Nucleus was stained with DAPI for 2 min at RT. Coverslips were mounted onto slides and complex formation was examined with an inverted Zeiss LSM 700 microscope.</u>

### Transfection of EHD4 plasmid and reverse co-immunoprecipitation

<u>hTERT-RPE1 cells were transfected using ESCORT IV Transfection Reagent (L 3287, Sigma-Aldrich), with pcDNA6.V5 and pcDNA6.V5 hEHD4 WT (kind gift from Dr. Markus Plomann, unpublished construct), according to manufacturer’s instructions: first, for 2 x 10e5 cells, 1.5 ug DNA and 2 ug ESCORT IV were diluted separately in OPTIMEM (Gibco), then mixed, and incubated 15 min at RT. After two washes in PBS, the mix was drop-wised on the cells, which were then incubated for 6 h at 37°C. The transfection was stopped by the addition of 20% FBS-DMEM/F12 medium and the cells were grown for 48 h at 37°C.</u>

<u>A co-immunoprocipitation was performed on the cells, using anti-V5 Agarose Affinity Gel antibody produced in mouse (A7345, MERCK) following the protocol described in the main methods section. EHD4 precipitation was confirmed using EHD4 antibody (1:500, ab153892 Abcam) and co-precipitated ARP-T1 was analyzed using ARP-T1 antiserum (1:2000, GP-SH6).</u>

## ACKNOWLEDGEMENTS

We are thankful to Prof. Rune Toftgård, to Prof. Anthony Oro<u>, and to Dr. Sergey Nikolaev</u> for their kind advice. We also acknowledge the Protein Analysis Facility of the University of Lausanne for the mass spectrometry assay. This study was funded by Swiss National Science Foundation (grant 310030-173102), “Fondation Professeur Placide Nicod” and “Fondation Dind Cottier pour la recherche sur la peau”.

## AUTHOR CONTRIBUTIONS

Please see corresponding file.

