## Extended data and supplementary table for "ARP-T1 is a ciliogenesis protein associated with a novel ciliopathy in inherited basal cell cancer, Bazex-Dupré-Christol Syndrome"

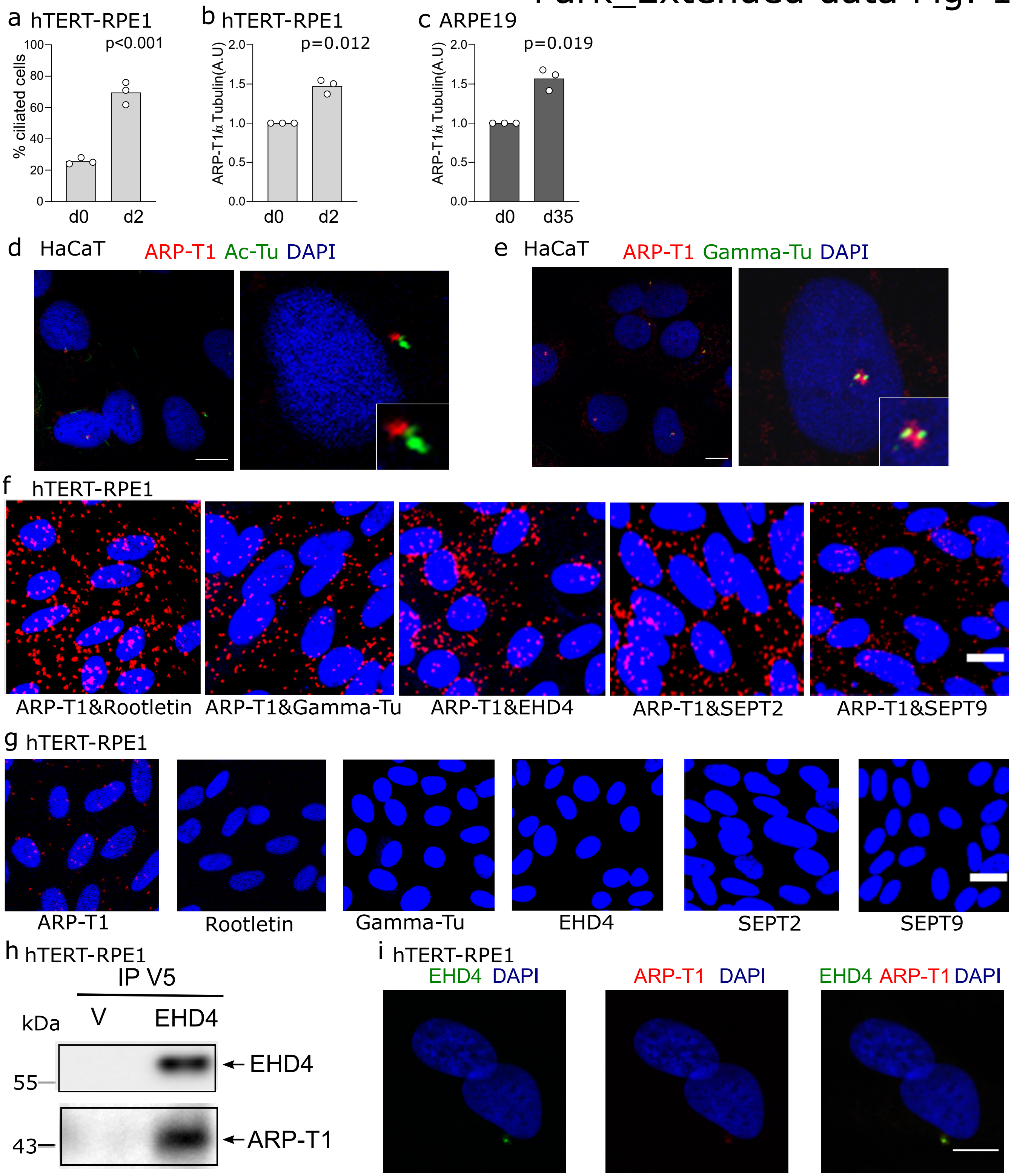


**Extended data Fig.1. Ciliogenesis and ARP-T1 increase after differentiation, and ARP-T1 localization and interactions.**

**a**, Percentage of ciliated hTERT-RPE1 cells under proliferating (d0) and differentiating (d2) conditions. Data are presented as means of the percentage. Each open circle represents one independent experiment (N=3). **b**,**c**, Quantification of ARP-T1 expression in hTERT-RPE1 (**b**) and ARPE19 (**c**). Data are presented as means of the fold change compared to the value of undifferentiated samples. Each open circle represents one independent experiment (N=3). **d**,**e**, Immunofluorescence stainings of ARP-T1 (red) and acetylated-tubulin (green) (**d**), and ARP-T1 (red) and gamma-tubulin (green) (**e**) in 7 days differentiated HaCaT cells. Nuclei are stained with DAPI (blue). Scale bar, 10 μm. **f**, Proximity-mediated ligation assays using ARP-T1 and rootletin, or gamma-tubulin, or EHD4, or septin 2, or septin 9 antibodies, in 48 h serum-starved hTERT-RPE1 cells. **g**, Proximity-mediated ligation assays using ARP-T1, rootletin, gamma-tubulin, EHD4, septin 2, or septin 9 antibody alone, in 48h serum-starved hTERT-RPE1 cells. **f**,**g**, Interactions are in red, nuclei are stained with DAPI (blue). Scale bar, 20 μm. **h**, hTERT-RPE1 cells were transfected with EHD4 and empty vector, serum-starved for 48 h, and immunoprecipitated (IP) with anti-V5 antibody-conjugated agarose, and analyzed by immunoblot with indicated antibodies. **i**, Immunofluorescence stainings of ARP-T1 (red) and EHD4 (green). Nuclei are stained with Hoechst (blue). Scale bar, 10 μm.

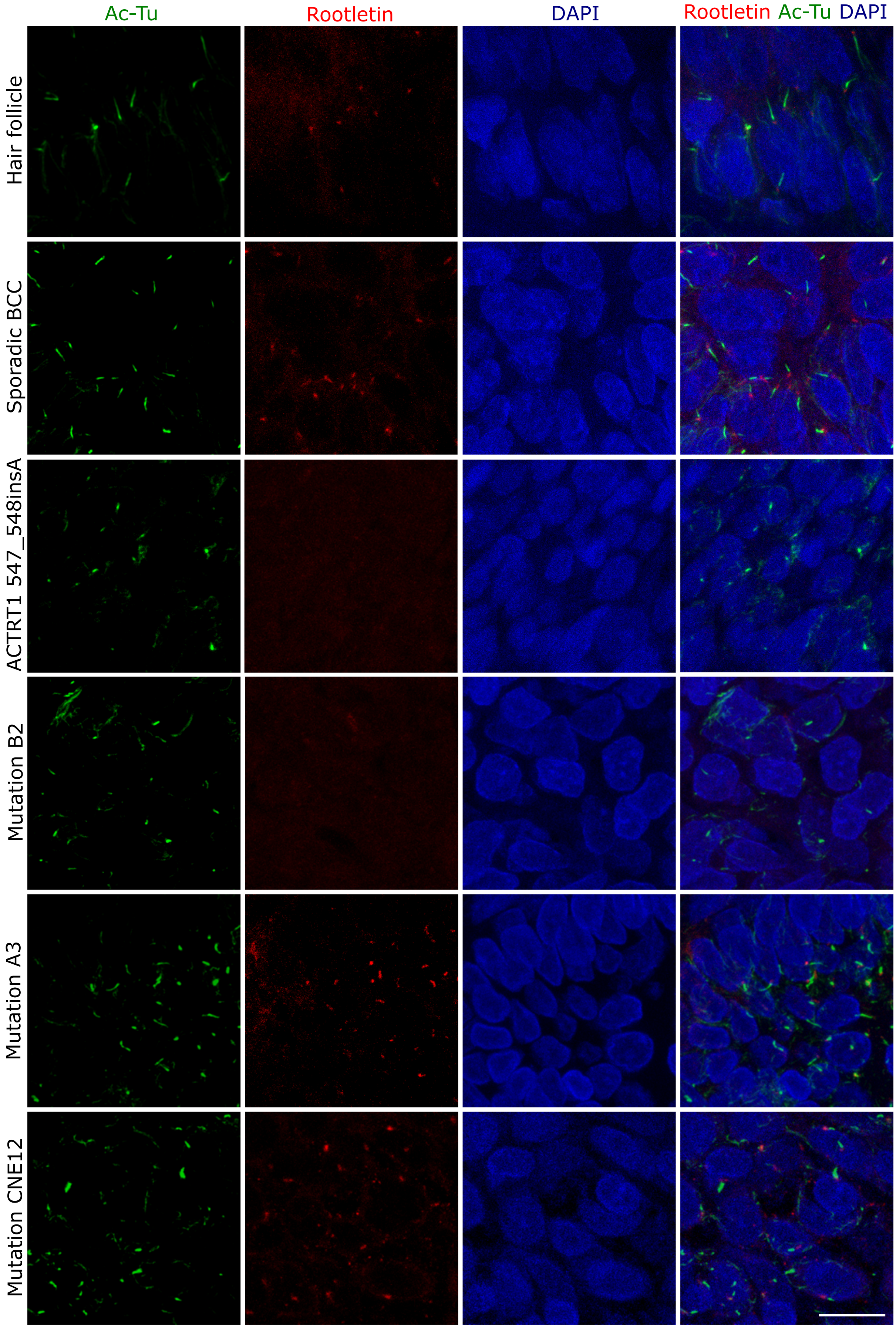


**Extended data Fig.2. Acetylated-tubulin and rootletin staining in patient samples.** Representative immunofluorescence images of acetylated-tubulin (green) and rootletin (red) in hair follicle, sporadic BCC and four BDCS (ACTRT1 547_548insA, Mutation B2, Mutation A3, Mutation CNE12). Cell nuclei are stained with DAPI (blue). Scale bar, 5 μm.

### Park_Extended data Fig. 3

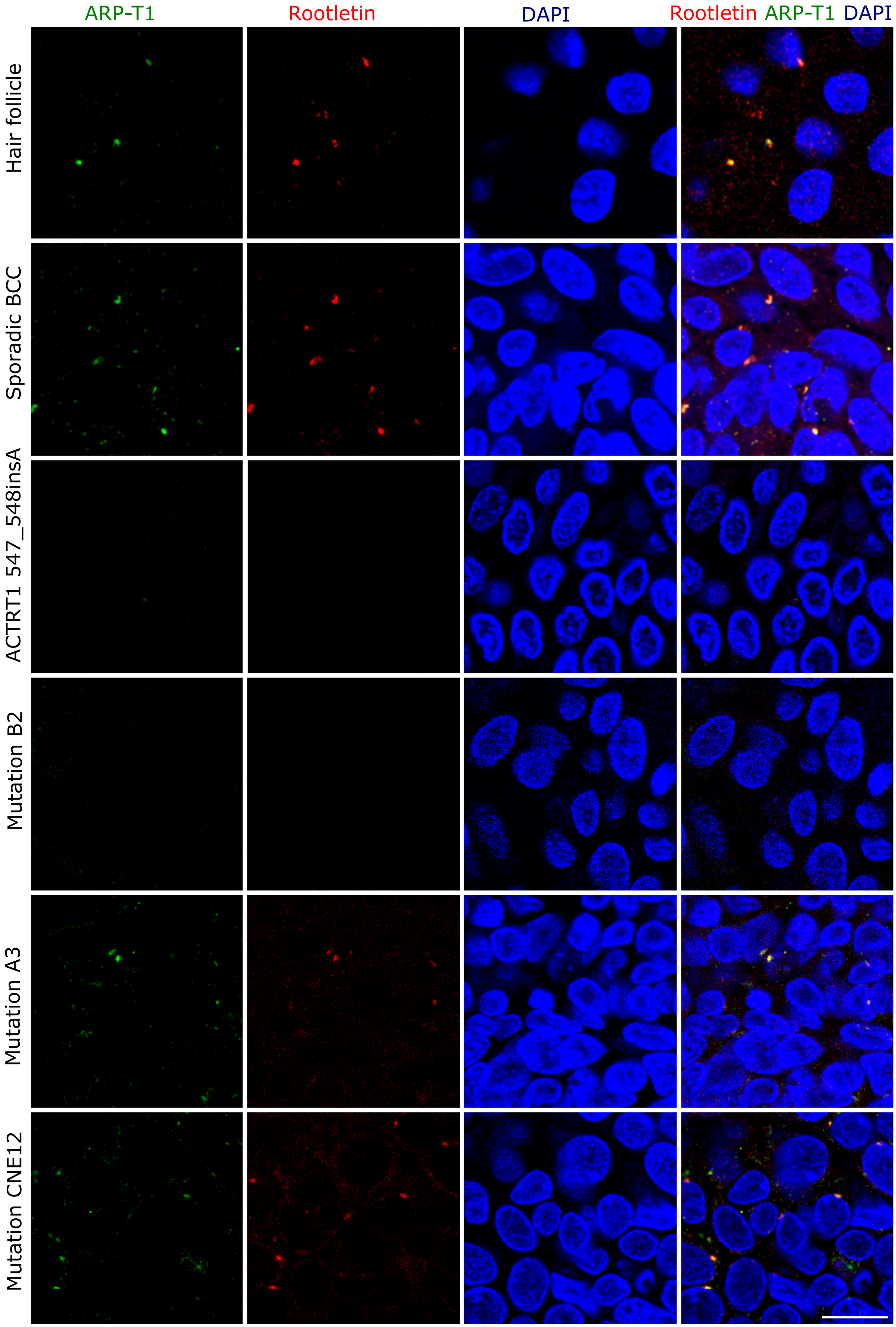


**Extended data Fig.3. ARP-T1 and rootletin staining in patient samples.**
Representative immunofluorescence images of ARP-T1 (green) and rootletin (red) in hair follicle, sporadic BCC and four BDCS (ACTRT1 547_548insA, Mutation B2, Mutation A3, Mutation CNE12). Cell nuclei are stained with DAPI (blue). Scale bar, 5 μm.

Park_Extended data Fig 4

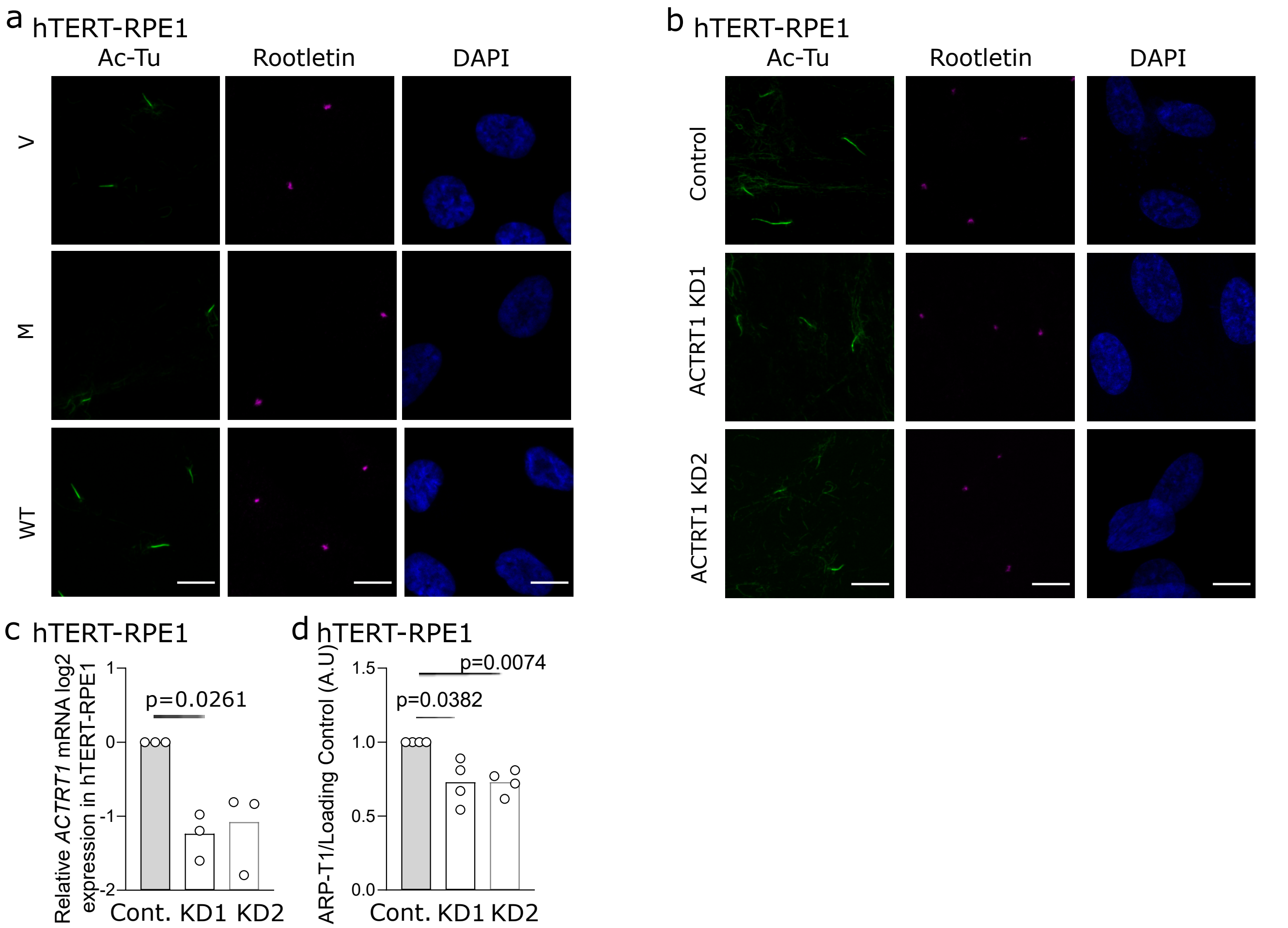


**Extended data Fig.4. ARP-T1 involvement in ciliogenesis in hTERT-RPE1 cells.**
**a**, Individual immunofluorescence stainings of acetylated-tubulin (green) and rootletin (pink) in 48 h serum-starved hTERT-RPE1 cells expressing an empty vector (V), or *ACTRT1* mutant (M), or *ACTRT1* WT (WT). Cell nuclei are stained with DAPI (blue). Scale bar, 10 μm. **b**, Individual immunofluorescence stainings of acetylated-tubulin (green) and rootletin (pink) in 48 h serum-starved control (Cont.) and *ACTRT1* KD hTERT-RPE1 cells. Cell nuclei are stained with DAPI (blue). Scale bar, 10 μm. **c**,**d**, Relative *ACTRT1* mRNA (**c**, N=3) and ARP-T1 (**d**, N=4) expression in control and *ACTRT1* KD hTERT-RPE1 cells. Data are presented as means of the fold change compared to the value of control cells. Each open circle represents one independent experiment (N=3).

Park_Extended data Fig 5

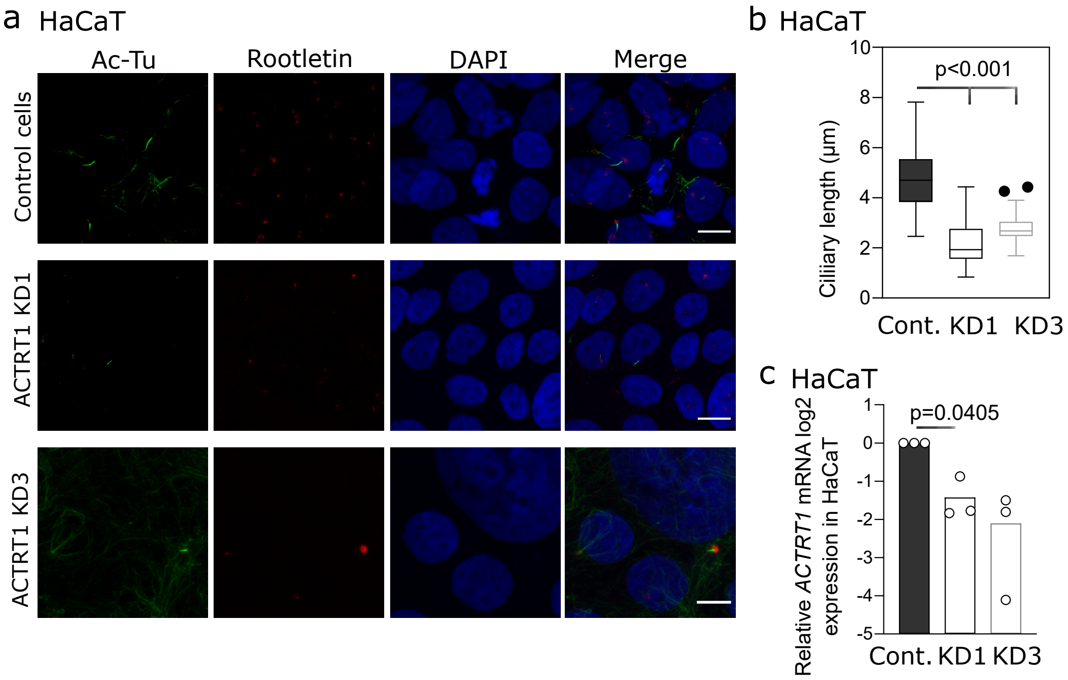


**Extended data Fig.5. ARP-T1 involvement in ciliogenesis in HaCaT cells.**

**a**, Immunofluorescence stainings of acetylated-tubulin (green) and rootletin (red) in 7 days differentiated control and *ACTRT1* KD HaCaT cells. Nuclei are stained with DAPI (blue). Scale bar, 10 µm. **b**,Quantification of ciliary length of **a**. Results are represented as Tukey box-plot. Black circles represent outliers. **c**, Relative *ACTRT1* mRNA expression in control (Cont.) and *ACTRT1* KD HaCaT cells. Data are presented as means of the fold change compared to the value of control cells. Each open circle represents one independent experiment (N=3).

### Park_Extended data Fig 6

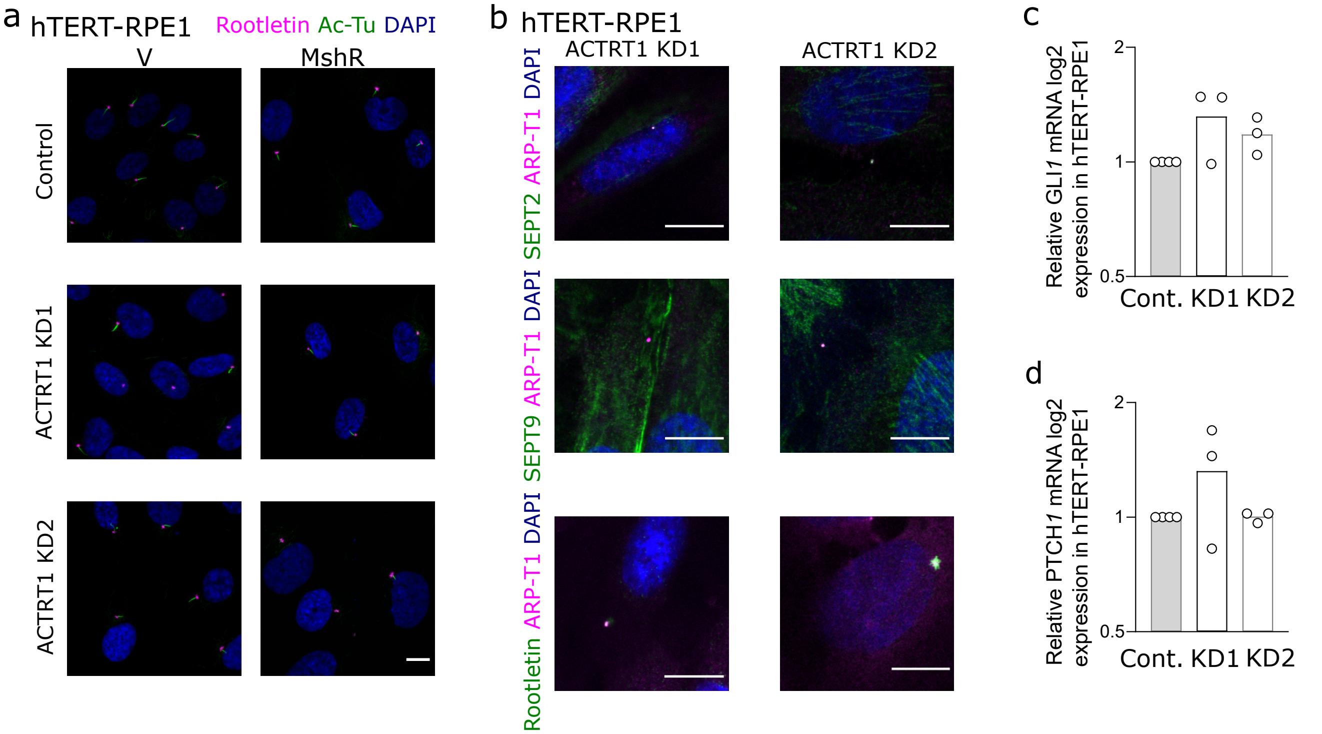


**Extended data Fig.6. ARP-T1 deficiency in hTERT-RPE1 cells.**

**a**, Immunofluorescence stainings of acetylated-tubulin (green) and rootletin (pink) in 48 h serum-starved hTERT-RPE1 cells expressing an empty vector (V) or *ACTRT1* mutant resistant shRNA (MshR). Cell nuclei are stained with DAPI (blue). Scale bar, 10 µm. **b**, Immunofluorescence stainings of ARP-T1 (pink) and septin 2 (green, top) or septin 9 (green, middle) or rootletin (green, bottom) in 48 h serum-starved control (Cont.) and *ACTRT1* KD hTERT-RPE1 cells. Cell nuclei are stained with DAPI (blue). Scale bar, 10 µm. **c**,**d**,Relative *GLI1* and *PTCH1* mRNA expression in control and *ACTRT1* KD hTERT-RPE1 cells. Data are presented as means of the fold change compared to the value of control cells. Each open circle represents one independent experiment (N=3).

| Pathway identifier | Pathway name | #Entities found | #Entities total | Entities ratio | Entities pValue | Entities FDR[a] |
|---|---|---|---|---|---|---|
| R-HSA-1430728 | Metabolism | 49 | 2135 | 1.91E-01 | 1.83E-03 | 9.14E-03 |
| R-HSA-392499 | Metabolism of proteins | 56 | 2012 | 1.80E-01 | 3.06E-06 | 1.69E-04 |
| **R-HSA-597592** | **Post-translational protein modification** | **32** | **1417** | **1.27E-01** | **1.65E-02** | **4.28E-02** |
| R-HSA-168249 | Innate Immune System | 31 | 1186 | 1.06E-01 | 2.32E-03 | 9.30E-03 |
| R-HSA-1643685 | Disease | 27 | 1173 | 1.05E-01 | 2.20E-02 | 4.40E-02 |
| **R-HSA-5653656** | **Vesicle-mediated transport** | **24** | **761** | **6.80E-02** | **6.64E-04** | **3.91E-03** |
| R-HSA-8953854 | Metabolism of RNA | 18 | 675 | 6.03E-02 | 1.66E-02 | 4.28E-02 |
| **R-HSA-199991** | **Membrane Trafficking** | **21** | **635** | **5.68E-02** | **8.15E-04** | **4.08E-03** |
| R-HSA-1640170 | Cell Cycle | 17 | 622 | 5.56E-02 | 1.59E-02 | 4.28E-02 |
| **R-HSA-422475** | **Axon guidance** | **20** | **558** | **4.99E-02** | **4.05E-04** | **2.43E-03** |
| R-HSA-168256 | Immune System | 169 | 2822 | 1.98E-01 | 9.33E-04 | 8.40E-03 |
| R-HSA-392499 | Metabolism of proteins | 157 | 2354 | 1.65E-01 | 8.16E-06 | 1.72E-04 |
| R-HSA-168249 | Innate Immune System | 86 | 1328 | 9.32E-02 | 0.00287 | 2.17E-02 |
| R-HSA-1280215 | Cytokine Signaling in Immune system | 86 | 1261 | 8.85E-02 | 6.58E-04 | 5.92E-03 |
| R-HSA-1280218 | Adaptive Immune System | 83 | 999 | 7.01E-02 | 9.83E-07 | 5.02E-05 |
| **R-HSA-5653656** | **Vesicle-mediated transport** | **63** | **824** | **5.78E-02** | **2.21E-04** | **2.43E-03** |
| R-HSA-8953854 | Metabolism of RNA | 62 | 782 | 5.49E-02 | 9.45E-05 | 1.23E-03 |
| **R-HSA-199991** | **Membrane Trafficking** | **59** | **665** | **4.67E-02** | **6.19E-06** | **1.42E-04** |
| R-HSA-8953897 | Cellular responses to external stimuli | 48 | 586 | 4.11E-02 | 2.89E-04 | 2.95E-03 |
| **R-HSA-422475** | **Axon guidance** | **50** | **584** | **4.10E-02** | **7.68E-05** | **9.98E-04** |

[a] FDR False Discovery Rate

**Supplementary Table 1:** Top10 of deregulated pathways in differentiated keratinocytes ACTRT1 WT vs MUT (top) and hTERT-RPE1 ACTRT1 WT vs MUT (bottom) analyzed with Reactome. Pathways in bold are linked to cilia and intracellular transport.
